# Machine learning models can identify individuals based on a resident oral bacteriophage family

**DOI:** 10.1101/2024.05.06.592821

**Authors:** Gita Mahmoudabadi, Kelsey Homyk, Adam Catching, Ana Mahmoudabadi, Helen Foley, Arbel D. Tadmor, Rob Phillips

## Abstract

Metagenomic studies have revolutionized the study of novel phages. However these studies trade depth of coverage for breadth. We show that the targeted sequencing of a small region of a phage terminase family can provide sufficient sequence diversity to serve as an individual-specific barcode or a “phageprint’’, defined as the relative abundance profile of the variants within a terminase family. By collecting ∼700 oral samples from ∼100 individuals living on multiple continents, we found a consistent trend wherein each individual harbors one or two dominant variants that coexist with numerous low-abundance variants. By tracking phageprints over the span of a month across ten individuals, we observed that phageprints were generally stable, and found instances of concordant temporal fluctuations of variants shared between partners. To quantify these patterns further, we built machine learning models that, with high precision and recall, distinguished individuals even when we eliminated the most abundant variants and further downsampled phageprints to 2% of the remaining variants. Except between partners, phageprints are dissimilar between individuals, and neither country-of-residence, genetics, diet nor cohabitation seem to play a role in the relatedness of phageprints across individuals. By sampling from six different oral sites, we were able to study the impact of millimeters to a few centimeters of separation on an individual’s phageprint and found that such limited spatial separation results in site-specific phageprints.

## Introduction

Viruses of bacteria, or phages, are among the most numerous and diverse biological entities on our planet. They play important roles as regulators of microbial ecosystems through rapid infection cycles and gene transfer events^1–3^. Yet, compared to their bacterial hosts, and despite their proven potential to transform fields such as medicine, agriculture and biotechnology^4,5,6–8^, phages remain as some of the least studied members of the human microbiome^9,10^. Even across familiar habitats such as the human body, the identity of phages and their corresponding bacterial hosts, their population structure, their modes of transfer between habitats, their co-evolutionary history with bacterial and human hosts, their role in health and disease, and other important topics remain relatively unexplored.

We chose to study phages residing in the human mouth as it represents a multifaceted and medically important ecosystem. Studies have revealed phages as highly abundant members of the human oral cavity, with distinct communities at sites of disease, capable of augmenting the bacterial arsenal of pathogenic genes^11–15^. These studies have relied on the shotgun metagenomic approach, in part because one of the defining features of viral genomes is the lack of a universally conserved sequence analogous to the 16S ribosomal RNA sequences in bacteria, which is used as a universal marker to draw conclusions about bacterial evolution and taxonomic classification^16,17^. This marker-based approach is indispensable to microbial ecology because it allows a high coverage depth of the 16S region, which in turn, enables precise and reproducible depictions of bacterial community compositions^18,19^.

Using current sequencing platforms, the trade-off for coverage depth is typically the coverage breadth (**SI Figure 1**). In comparison to the marker-based approach, shotgun metagenomics provides much greater breadth of coverage and offers several advantages. However, it suffers from several key disadvantages. The coverage depth is often heterogeneous and remains comparatively low in these studies, meaning that the *de novo* assembly of genomes from complex environments remains a significant challenge^20,21^, even for abundant members with relatively short genome lengths^22,23^. Moreover, the genomes assembled through shotgun metagenomics are often consensus genomes or an average representation of similar genomes within an environment^24^.

Due to these technical challenges, the marker-based approach allows orders of magnitude greater coverage depth by focusing the reads on a small genomic segment, and thus provides a much higher resolution view of microbial communities. The targeted approach is therefore widely used to complement shotgun metagenomic depictions of bacterial communities^25,26^. Because of their high mutation rates and rapid turnovers, viral genomes are incredibly diverse, and the study of the sequence diversity within a virus family could be much more deeply explored through targeted sequencing. Even within a single “species”, viral genomes exist as a collection of related variants, which are often described as “quasispecies’’ or as a “mutant spectrum”. The mutant spectra of RNA viruses is well described in early and recent studies of RNA phages and RNA viruses, particularly for lab strains^27–30^. DNA phages, on the other hand, are less studied within this framework, primarily because they have lower mutation rates compared to RNA phages^31^. Even less explored are the mutant spectra of DNA phages within a dynamic host environment.

As such, the overarching aim of this study was to apply targeted sequencing to understudied DNA phages in their native context, to explore their inter-and intra-personal diversity, their spatial patterns of distribution, as well as temporal dynamics in a large-scale and high-resolution fashion that allows for observing their individual variants as well as the collective mutant spectra. Thus, we first had to choose regions within phage genomes on which to perform targeted sequencing. While one could relatively easily target sequences of well characterized phages, we were motivated to create a roadmap for mining metagenomic datasets and shedding light on understudied phages.

Towards this goal, we first developed and benchmarked Metagenomic Clustering by Reference Library or MCRL, which is an algorithm for the identification of non-redundant gene families within a metagenome^32^. In a previous study, we then applied MCRL to oral metagenomes of seven individuals from two studies conducted in two different continents^33,34,35^. By focusing the search on the terminase (large subunit) gene families, we were able to narrow down the search from thousands of viral gene families to seven non-homologous terminase families that were shared across individuals in these two studies^35^.

In the absence of a genomic taxonomy for viruses, we have referred to those phages that encode members of the same terminase family as members of the same phage family^35^. This notation is predicated on previous studies, including our own^36^, that have shown no significant sequence similarity between terminase sequences of unrelated phages^37,38^ as well as studies that have used the terminases to build phage phylogenetic trees^39,40^. Moreover, we focused our search on terminases because they are among the most functionally-conserved genes in double-stranded DNA phage genomes^41,42^. Unlike several other viral genes such as integrases and lysins, terminases lack bacterial homologs, and thus, are considered to be unique to phages^43^. Additionally, we have previously successfully used terminases to probe phage-bacteria interactions within a complex host environment, namely the termite gut^44^.

To test whether we were successful in identifying terminase families that were prevalent enough in the human phageome to be practical experimental targets, we searched for them across hundreds of metagenomic samples from the Human Microbiome Project (HMP)^45^ spanning ∼100 individuals and 18 body sites^35^. Remarkably, we showed that despite the individual-specific nature of the human virome and the small number of individuals from which these terminase families were originally identified, they are prevalent across the HMP cohort. In this study we chose to focus on HB1 and HA terminase families as they were the two most prevalent families, detected in most individuals within the HMP cohort^35^. In the following paragraphs we summarize some of our earlier findings, particularly those pertinent to HA and HB1 terminase families.

To identify the putative habitats of the phages encoding these terminase families, we searched through ∼4000 environmental metagenomes from the IMG/VR^46^ and IMG/M^47^ databases comprising numerous distinct habitats, in addition to ∼100 environmental metagenomes from the VIROME database^48^. Most terminase families were found to be largely human-associated, and instances where remote homologs were found in environmental phages, the human-derived phage sequences were phylogenetically distinguishable from their environmental counterparts. Additionally, by examining various body sites, we showed that most terminase families were primarily localized to the human oral cavity. The HB1 terminase family was found as an exception given that it is detected also in the human gut, though we showed that the oral and the gut-derived HB1 terminase family members were phylogenetically distinct.

Through experiments where we separated the bacterial and viral fractions of oral samples, we were able to demonstrate that the HA phage family is likely lysogenic and infects various species of the *Steptococcus* genus, whereas the oral HB1 phage family is likely lytic, and its host species remains to be discovered. Moreover, we show the positions of the closest HA and HB1 terminase homologs in previously sequenced full phage genomes **(SI Figure 2**). Additionally, through selection pressure analysis and alignment of functional motifs, we showed that HA- and HB1-encoding phages are likely functionally active members of the human oral virome. Finally, we designed primers to target these phage families using their respective terminase families within oral samples from nine individuals and showed that we could indeed reliably capture them experimentally. The primers for HA and HB1 are provided again in this study (**SI Table 1**).

In this study, we target the HA and HB1 terminase families to obtain at least several thousand sequences per terminase family, per oral sample, and thereby increase the resolution or the coverage depth by several orders of magnitude from our previous study. By creating instructional videos and collection kits, we enabled citizen scientists to gather ∼700 samples spanning ∼100 individuals residing in different parts of the world (**Figure 1**). We will demonstrate that at high resolution, the mutant spectrum derived from members of just a single phage terminase family can already serve as a fingerprint, or a “phageprint” – highly unique to an individual. Phageprints were not observable through our earlier study of metagenomic datasets^35^, and demonstrate the power of combining metagenomic mining with targeted sequencing to put a spotlight on uncharacterized phage families and their sequence diversity in their native contexts.

**Figure 1.**
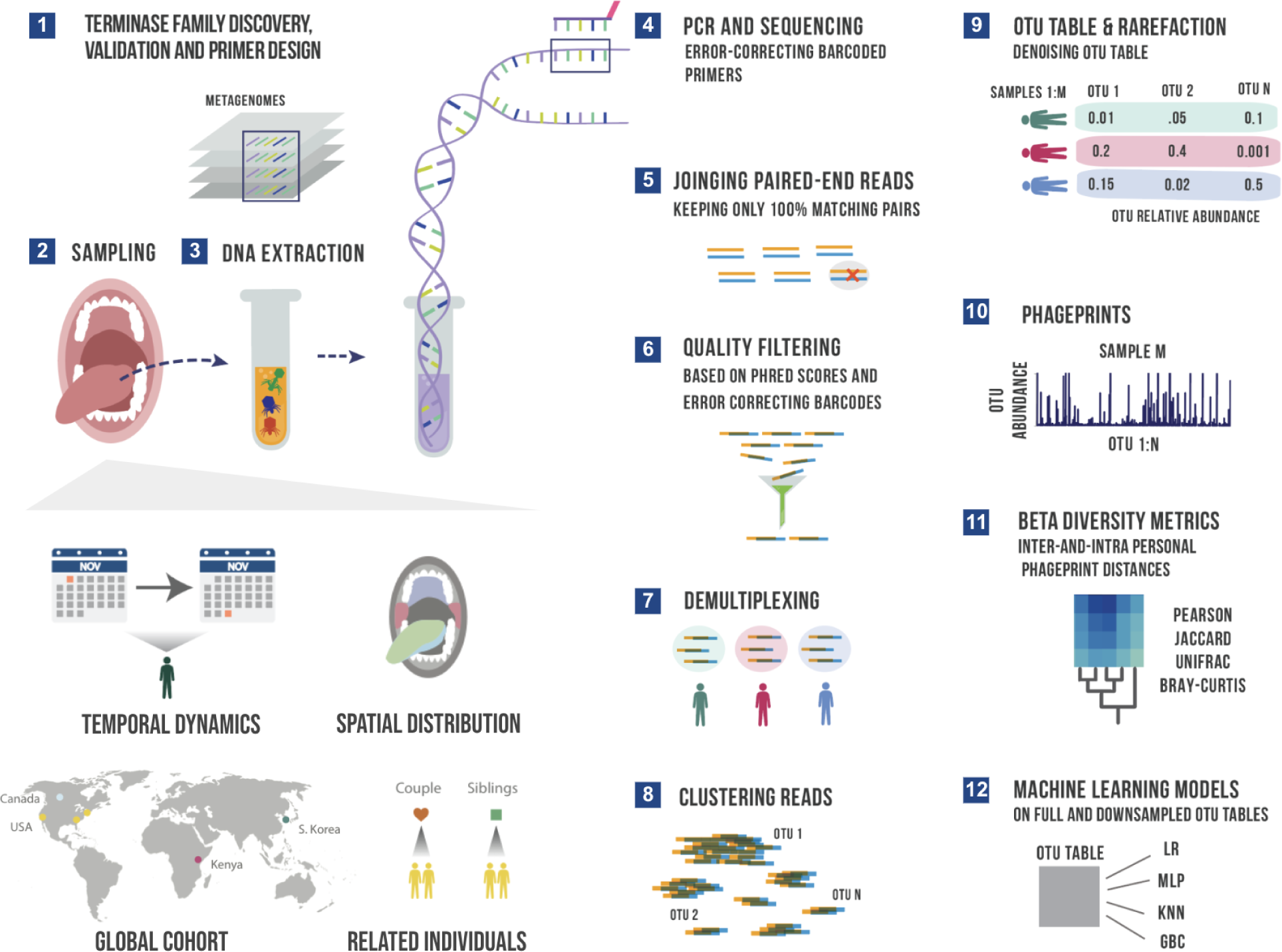
A schematic summary of the main experimental and bioinformatic methods: 1) Discovery of ubiquitous phage families by examining large terminase sequences that occur across different metagenomic datasets described in our earlier work^35^, 2) experimental sampling of several cohorts for temporal and spatial analysis of phageprints in related in unrelated individuals, 3) DNA extraction from oral biofilm samples, 4) PCR using barcoded primers followed by PCR clean-up and paired-end sequencing, 5) joining paired-end reads to eliminate sequencing errors, 6) additional quality control steps to further eliminate errors based on Phred scores and error-correcting barcodes, 7) demultiplexing of reads based on their barcode sequence and linking sequences to the sample they originate from, 8) gathering reads from all samples and clustering them based on sequence similarity into Operational Taxonomic Units (OTUs), 9) counting the number of sequences belonging to each OTU from each sample (i.e. constructing an OTU table), and rarefying the table so that each sample is represented by the same total number of sequences, and denoising the OTU table to eliminate OTUs with relative abundances below an experimentally determined reproducibility threshold, 10) visualizing phageprints which are the relative abundance profiles of OTUs (1 through N) in a given sample, 11) performing various downstream diversity analysis using the constructed OTU table as the basis, 12) creating machine learning models based on full and downsampled OTU tables. These model types include Logistic Regression (LR), Multi-Layer Perceptron (MLP), K-nearest Neighbor (KNN) and Gradient Boosting Classifier (GBC). Note that these steps are performed separately for HA and HB1 sequences.

By examining phage terminase families at 6 different oral sites, and by comparing phageprints of individuals living across the globe, we were able to study the effect of spatial separation, ranging from several millimeters to thousands of kilometers. We found that the spatial separation of just a few centimeters - the distance between an individual’s gingival sites and the hard palate, for example - already results in highly distinct phageprints for the HA phage family. In contrast, HB1 phageprints from different oral sites within an individual were highly similar. Additionally, we found that neither genetics nor cohabitation seem to play a role in the relatedness of phageprints across individuals.

Furthermore, by daily sampling of phageprints from the tongue dorsum over the course of a month across ten individuals we continued to see individual-specific phageprints with many variants that persisted over time. We also identified variants that were fluctuating concordantly in partners. Through various diversity metrics we quantified the inter-and intra-personal distances between phageprints as a function of space and time. We used machine learning models to further quantify the identifiability of an individual’s phageprint and showed remarkably high model performances on unseen data. These models had very high performances even as the most abundant variants were removed and even when 98% of the remaining variants were randomly removed.

## Results

### Humans harbor diverse, personal phageprints that are persistent in time

From a methodological standpoint, targeted sequencing of teminase families is very similar to 16S sequencing^18,49^. Using barcoded primers, we employed PCR and next generation sequencing to attain millions of paired-end reads for each terminase family (**Figure 1**). We took stringent measures against contaminants by 1) conducting our DNA extraction, PCR and post-PCR experiments in separate physical spaces, and 2) running five no template control reactions for every PCR run, as well as three no-sample DNA extraction reactions for every DNA extraction run to ensure there are no contaminants in the DNA extraction kits. Upon sequencing and performing several quality control filters, the reads were demultiplexed based on their barcoded primer sequence. Using error-correcting DNA barcodes, we were able to detect errors and removed sequences if they contained errors in their barcode. Furthermore, we eliminated nearly all sequencing errors by using paired-end reads which covered the full length of both terminase families (300 bp) and allowed only paired sequences with 100% match across the entire sequence (see **Materials and Methods**).

All reads derived from the same terminase family were then pooled and clustered based on their DNA sequence similarity into Operational Taxonomic Units (OTUs), or what we will interchangeably refer to as variants. An OTU table is constructed wherein the number of reads belonging to each OTU (columns) within each sample (rows) is denoted. Using the OTU table, we can plot the relative abundances of each OTU within a sample. As a shorthand, we refer to this plot as a phageprint.

With bacterial 16S data, sequences are generally clustered at 97% sequence similarity into OTUs. At this threshold, each OTU is conventionally referred to as a bacterial species. In the absence of convention for handling viral targeted sequencing data, we have used here various sequence similarity thresholds for clustering including 100% sequence similarity, thereby allowing only identical sequences in each cluster. We found the results to be largely robust to variations in the sequence similarity threshold (see Materials and Methods: Examining the effect of OTU sequence similarity threshold, **SI Figure 3**).

As an example, we show the HA phageprint from a subject’s tongue dorsum (top surface) at two time points (**Figure 2.a**). As shown in this figure, and across all other phageprints we have constructed for both terminase families, each phageprint is dominated by a small number of variants or OTUs (typically one or two). In addition to these OTUs, there are many OTUs with abundance values that are low but reproducible, and some that are fairly persistent in time within each subject. Generally, the dominant OTUs are not the same across different subjects.

**Figure 2.**
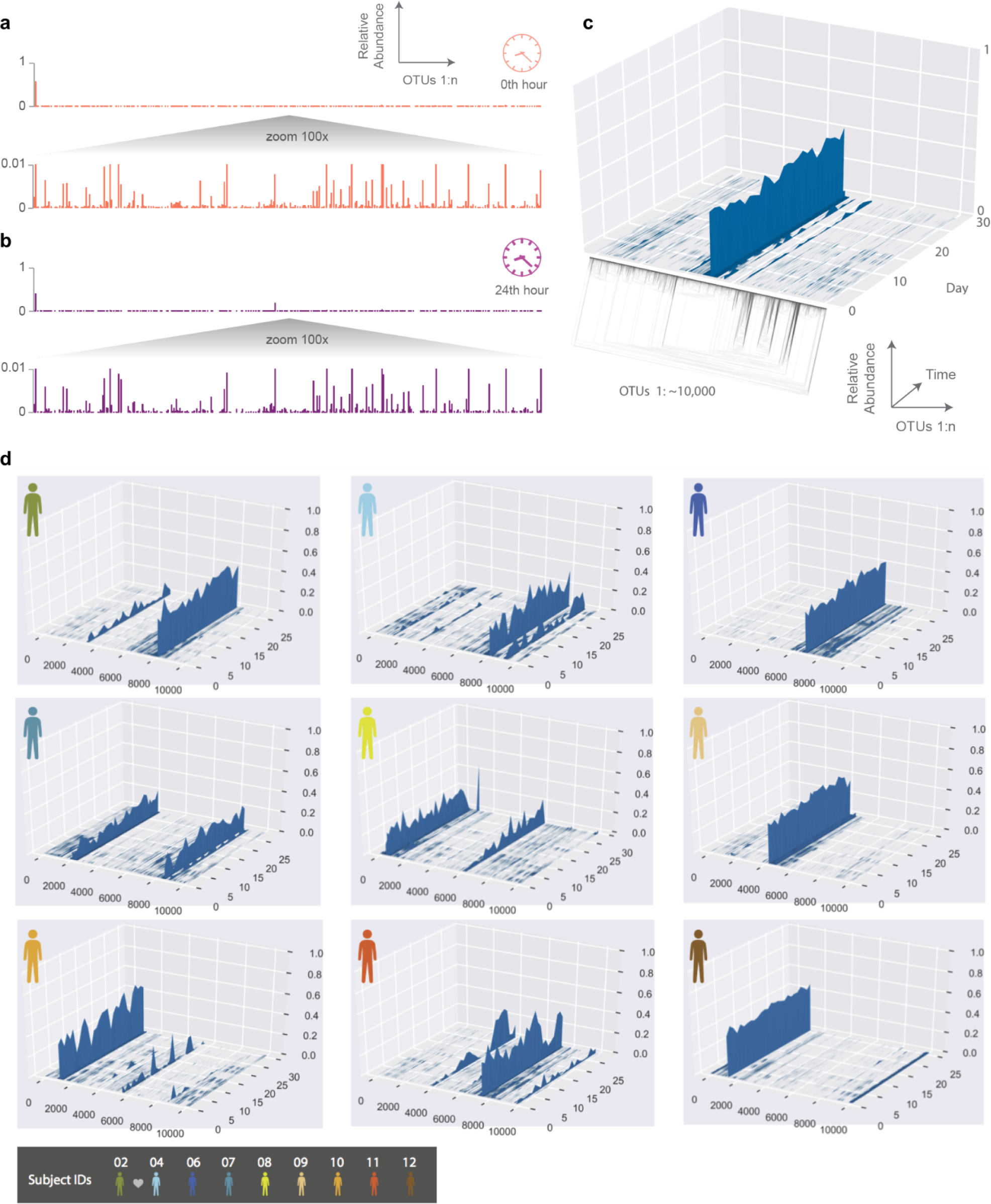
The temporal dynamics of an individual’s phageprint over the course of a month (on average 25 daily samples were collected during this period). a-b) HA phageprints from subject 37 at two different time points, (a) 0th time point, right after brushing tongue dorsal and teeth surfaces and b) 24 hours after the initial time point (no brushing in between time points). Each phageprint is derived from the analysis of 4000 sequences. OTUs are defined at 98% sequence similarity. c) HB1 phageprint temporal dynamics on subject 1’s tongue dorsum. The x-axis contains OTUs ordered according to the depicted phylogenetic tree of the OTU sequences (the phylogenetic tree is provided largely to serve as a schematic). Each OTU is composed of identical sequences (i.e. 100% sequence similarity threshold). The y-axis depicts the relative abundance of each OTU, and the z-axis shows the fluctuations in relative abundance of each OTU in time. d) Depictions of HB1 phageprint temporal dynamics in different subjects. The format of these plots is the same as that panel c, and the order of OTUs is based on their phylogenetic distance and identical across all plots. All samples are collected from the tongue dorsum. Note that subject 2 and 4 are partners, and their phageprints share some main features.

Before probing a larger number of individuals, we aimed to quantify our pipeline’s detection and reproducibility thresholds to understand what levels of OTU temporal fluctuation is biological versus technical. To that end, we obtained 3 different samples from a subject’s tongue dorsum. We then performed DNA extraction and PCR separately on each sample and sequenced these samples. The logic behind this experiment was to capture a lumped measure of noise arising from various experimental processes depicted in **SI Figure 4**. We show that the relative abundance of the variants making up each phageprint across these three samples are highly reproducible, and the maximum standard deviation for OTU relative abundances was less than 0.007, with the majority less than 0.002 and close to 0 (**SI Figure 5**). Moreover, we flagged OTUs that had appeared in only one or two samples out of three. As expected, we observed that the number of reproducible OTUs increases as a function of the relative abundance threshold, and all OTUs with greater than 0.001 relative abundance were reproducible across all three samples (**SI Figure 6**). Thus, we arrived at 0.001 relative abundance as the reproducibility threshold for OTUs, and denoised OTU tables by eliminating OTUs that did not meet this threshold across any of the samples. We have performed similar benchmarking studies on a larger number of subjects and included separate sequencing runs to account for any variation that may be introduced by a sequencing run (**SI Figure 7**). In short, through stringent quality control filters and benchmarking of our experimental and bioinformatic workflow, we showed that phageprints are highly reproducible (**see Materials and Methods**).

To further explore the temporal dynamics of these phageprints, ten subjects collected biofilm from the tongue dorsum every 24 hours for a month though on average subjects returned samples from 25 days as they missed to sample some days. The HB1 phageprint temporal dynamics on a subject’s tongue dorsum is depicted in **Figure 2**. Here, to provide a more detailed view, we cluster the HB1 sequences into OTUs based on 100% sequence similarity, or in other words, we are depicting the relative abundance of individual sequences.

Given the dynamic nature of an ecosystem like the human mouth, it is counter-intuitive that over a month, the main features of each phageprint is preserved in all subjects. However, as we will investigate further, there are fluctuations that are biological rather than technical. A global trend is that the dominant OTUs typically remain dominant throughout the sampling period in all subjects (**Figure 2**). This observation is especially interesting in light of the inter-and intra-personal differences in diet and oral hygiene practices which subjects reported on (**SI Figure 8**).

To make quantitative pairwise comparisons between phageprints we employed several commonly used metrics such as Bray-Curtis and Unifrac, and in doing so, we distill the comparison of thousands of sequences from any two samples to a single score. All distance metrics paint similar pictures of the HB1 terminase family, depicting it as highly individual-specific and persistent in time (**SI Figure 9**, **Figure 3**). Because phageprints in different individuals have such distinct compositions, abundance-based metrics are especially suitable for describing them. However, even the binary Jaccard distance metric which does not consider variant abundances point to a similar conclusion. As is expected from the heat maps shown in **SI Figure 9**, the intra-personal distances are markedly lower than the inter-personal, with the notable exception being subjects 2 and 4, who are partners (**Figure 3**).

**Figure 3.**
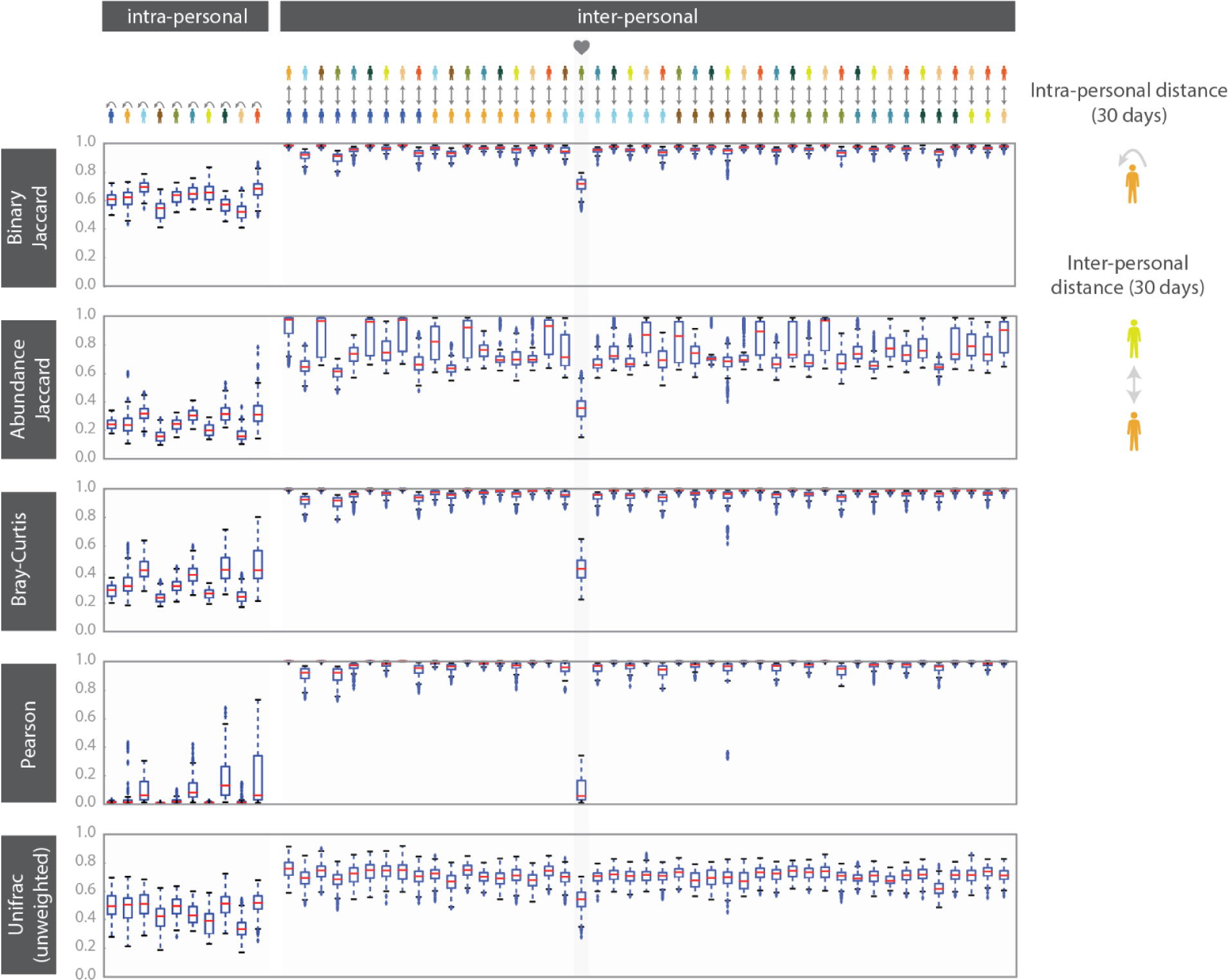
HB1 phageprint temporal dynamics quantified using pairwise distance metrics and visualized using a) heatmaps and b) box-and-whisker plots. The pairwise distance metrics include: Peason distance (1-Pearson correlation), Binary Jaccard, Abundance Jaccard, Bray Curtis and unweighted Unifrac. a) The heatmap scale applies to all heatmaps shown. Subjects 02 and 04 are partners. Samples from each subject are chronologically ordered. b) Intra-and inter-personal distances between HB1 phageprints in 10 subjects, over the span of a month. The outliers defined as those outside of the 1.5 x IQR (inter-quartile range) are denoted by “+”. The box-plots corresponding to the comparisons between the couple in this study are highlighted.

### Machine learning models detect with high precision and recall an individual’s phageprint even when phageprints are heavily downsampled

In addition to these distance metrics, we were motivated to build machine learning models whose performance could further quantify the predictability of an individual’s phageprint within the temporal cohort. We first built several types of machine learning models, including Logistic Regression (LR), K-Nearest Neighbor (KNN), Gradient Boosting (GBC), and Multi-Layer Perceptron (MLP), each of which perform a binary classification of an individual’s phageprint from the rest (i.e. one-versus-rest models). The input to these models was the OTU table, where the rows are samples (i.e. day 1 to 30 for each subject) and the columns are the OTUs. Across the temporal cohort consisting of ten individuals, ∼7300 HB1 OTUs were collectively detected. This table was split for training (70%) and testing (30%) such that models would be trained on 70% of the time points from each individual. To quantify the performance of the models, we performed ten iterations of random train/test splits and report the median and the 95% confidence intervals for the Area Under the Precision-Recall curve (AUPR) and the Area under the Receiver-Operator Curve (AUROC).

All model types performed remarkably well with very high performances for both the Logistic Regression and the Multi-Layer Perceptron model types (**Figure 4**, **SI Table 2**). We performed the same exercise on an OTU table built from HA terminase family OTUs, and arrived at similarly high model performances (**SI Figures 10-11, SI Tables 4-5)**. It is important to note that we excluded subject 4 from this particular analysis because we wanted to measure the model’s performance for unrelated individuals, as partners’ coevolving phageprints would be a confounding factor. We chose to build Logistic Regression models moving forward since this model type performed very well for both phage terminase families.

**Figure 4.**
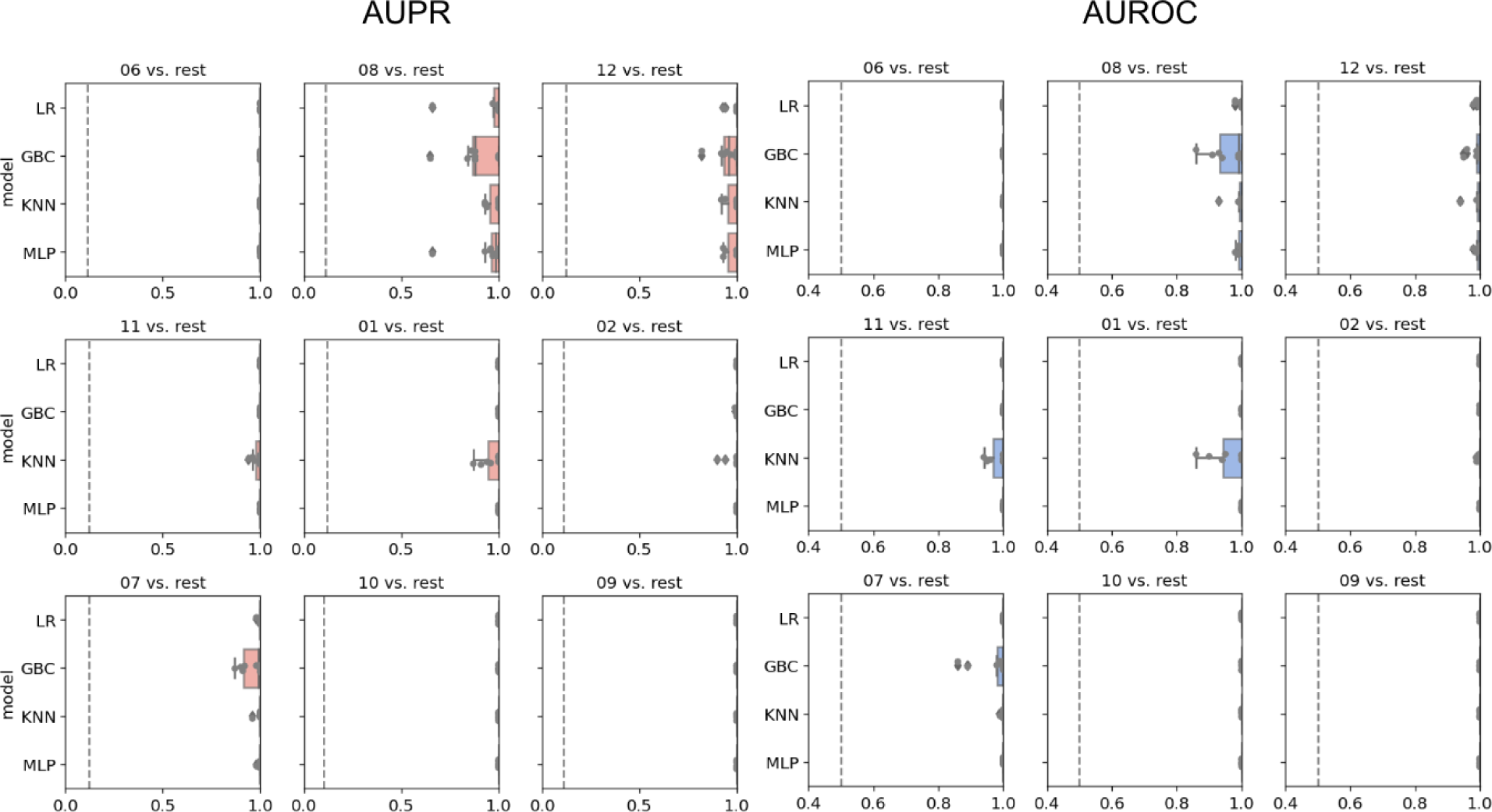
Machine learning one-versus-rest models built to distinguish between one person’s HB1 phageprints from the rest. Salmon and blue panels represent boxplots of the Area Under the Precision Recall (AUPR) Curve and the Area Under the Receiver Operator Curve (AUROC) values, respectively. The null values are shown as dashed lines, which for AUROC is equal to 0.5 and for AUPR is equal to the prevalence of the positive class. The four model types shown on the y axes are Logistic Regression (LR), Gradient Boosting Classifier (GBC), K-Nearest Neighbor (KNN), and Multi-Layer Perceptron (MLP). For each model type 10 models are built based on 10 different splits of the data into training and testing portions. Subject IDs are shown at the top of each panel, such that “06 vs. rest” for instance, corresponds to model performances on test data distinguishing subject 6 phageprints from all other subjects’ phageprints.

Given that phageprints are dominated by one or two OTUs, it is reasonable to assume that the exclusion of these dominant OTUs would dissolve the individual-specific and time-persistent nature of phageprints. To formally test this assumption, we removed the top ten most abundant OTUs of each sample from the entire dataset. A total of ∼600 OTUs were removed from the dataset, removing on average two thirds of the reads from each sample. Upon removing these OTUs, we rescaled the OTU table such that the relative abundance of the remaining OTUs would again add up to 1. To our surprise, the exclusion of the top most abundant OTUs still resulted in nearly perfect classification (**SI Tables 6-7**). We further randomly downsampled to 2% of the total remaining OTUs, resulting in just 226 OTUs, and rescaled the resulting OTU table as previously described. The performance of the models still remained nearly as high as before (**SI tables 8-9**).

The reason for the repeated observation of phageprints even when drastically subsampled, is due to the fact that many low-abundance OTUs have individual-specific patterns of occurrence. By hierarchical clustering of this small subset of the original OTU table (**SI Figure 12**), most samples from the same individual cluster together, and thus, machine learning models can easily pick out an individual’s phageprint from others even using a small fraction of the total data for each subject.

### Less than 1% of OTUs are shared across all subjects

We measured the sharing of OTUs across subjects by collapsing the OTU table into a table of subjects by OTUs rather than samples by OTUs, such that if an OTU was identified at any point within the sampling period (∼30 days), it is given a value of 1, and 0 otherwise. With this binary table, we created an UpSet plot where the number of OTUs unique to each subject as well as the number of OTUs shared between different sets of subjects is shown (**SI Figure 13**).

Less than 1% (∼0.8%) of all OTUs were detected across all subjects. The relative abundance of these generalist OTUs per subject is hierarchically clustered and shown in **SI Figure 14**. Again, we see that partners cluster most closely together even based on this small subset of OTUs. Finally, a much higher percentage of total OTUs, about 85%, are detected in at least two subjects, and the rest are only detected in one individual. Based on these results, we can conclude that while the same variants may appear in different subjects, the individual specificity of phageprints emerge in large part because the relative abundances of variants is often individual-specific.

### While phageprints are largely persistent in time, they are temporally dynamic with partners sharing some concordantly fluctuating OTUs

To further examine the temporal aspect of this data, we plotted the number of days that each OTU was detected within each subject (**SI Figure 15**, **SI Table 10**). The average number of days an OTU was detected in this temporal study ranged from 5 to 7 days depending on the subject, though there were many OTUs within each subject that appeared across nearly all time points. We dubbed OTUs detected in at least 20 days as highly persistent OTUs. This threshold was chosen because it represents two standard deviations from the mean number of days an OTU is detected, averaged across all subjects (**SI Table 10**).

To further quantify the temporal dynamics of OTUs in each subject, we measured the coefficient of variation for persistent OTUs (**SI Figure 16**), and selected outliers, which represent highly dynamic yet persistent OTUs for each individual. **SI Figure 17**, shows some representative plots of such outlier OTUs, and demonstrates these fluctuations. Moreover, we examined the OTUs that are shared between partners (subjects 2 and 4) and plotted their relative abundance. Interestingly, we discovered that certain OTUs are concordant in their temporal fluctuations in these two individuals. We demonstrate some representative plots for such OTUs (**SI Figure 18**). We also found OTUs whose dynamics are very similar across most time points except for a few when one partner’s OTU rises by several folds and falls back down (**SI Figure 19**). Future studies are needed to shed light on the underlying mechanisms giving rise to such fluctuations.

### Comparative analysis of phageprints across siblings, couples, and unrelated individuals residing on different continents

Given the ubiquitous presence of HA and HB1 in both our cohort and in metagenomic datasets we wondered whether subjects residing in the same country might have more similar phageprints. Neither from abundance-based nor phylogenetic distance comparisons did we find an indication that individuals residing in the same city or country share more similar phageprints (**Figure 5**). Instead, we continued to find that individuals typically have highly unique phageprints.

**Figure 5.**
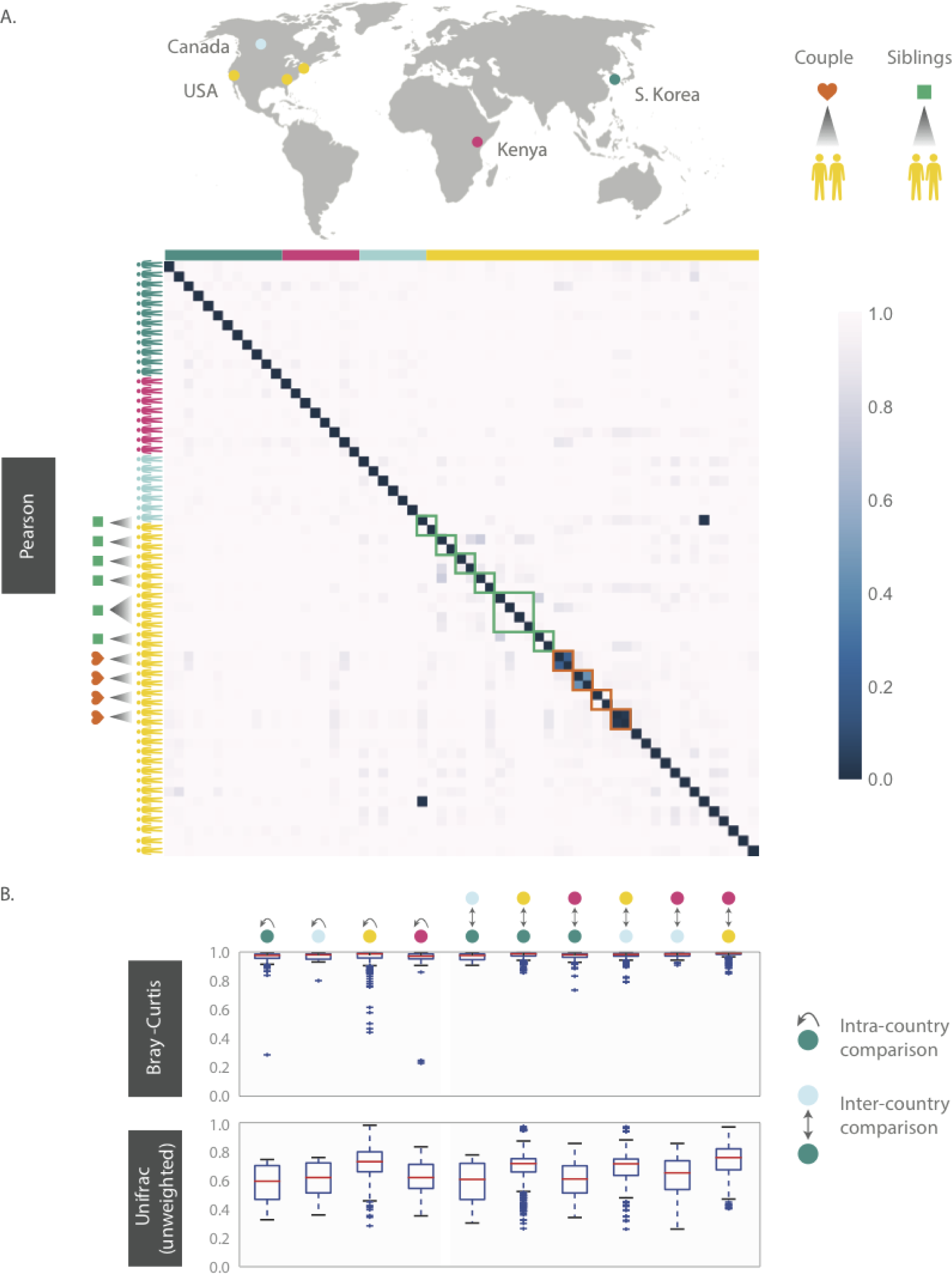
HB1 phageprints across 61 individuals residing across different parts of the globe. Samples are obtained from the tongue dorsum. A) Pearson distance (1 – Pearson correlation) is shown as a heatmap. A subset of individuals residing in the U.S. are either partners or siblings. Green and red boxes are drawn around samples from each sibling group and partners, respectively. B) Intra- and inter-country distances from pairwise comparisons made using Bray-Curtis and unweighted Unifrac distance metrics. The outliers are denoted as points outside of the 1.5 x IQR (interquartile range). Related individuals are excluded from this analysis.

Even siblings who were either living in the same household or had previously, do not have any more similar phageprints than unrelated individuals. In fact, one of the four sibling groups is identical twins (**Figure 5**). However, 3 out of 4 partners in this study exhibited highly similar phageprints (**Figure 5**). The dissimilar couple may be due to celiac disease diagnosed in one of the partners, which is known to alter oral ecology^50^. These results suggest that genetics and cohabitation do not significantly impact a person’s oral phageprint. Thus, we suspect phageprints of partners coevolve while phageprints of even cohabiting individuals evolve independently through time. To further test these trends, larger studies encompassing a greater number of individuals and regions in the world are needed.

### Within the same individual there are site-specific phageprints

Thus far, all phageprints shown are those sampled from the tongue dorsum. In order to examine the spatial patterns of phageprints we obtained additional oral samples covering 9 individuals and 6 oral sites (courtesy of Bik *et al*.^51^). **Figure 6** shows the HB1 phageprints of a subject at four oral sites where the HB1 terminase family was found. Clearly, different oral sites in this subject have very similar HB1 phageprints. When examining all HB1 positive samples, an immediately recognizable pattern is that the HB1 phageprints from different oral sites within an individual are highly correlated.

**Figure 6.**
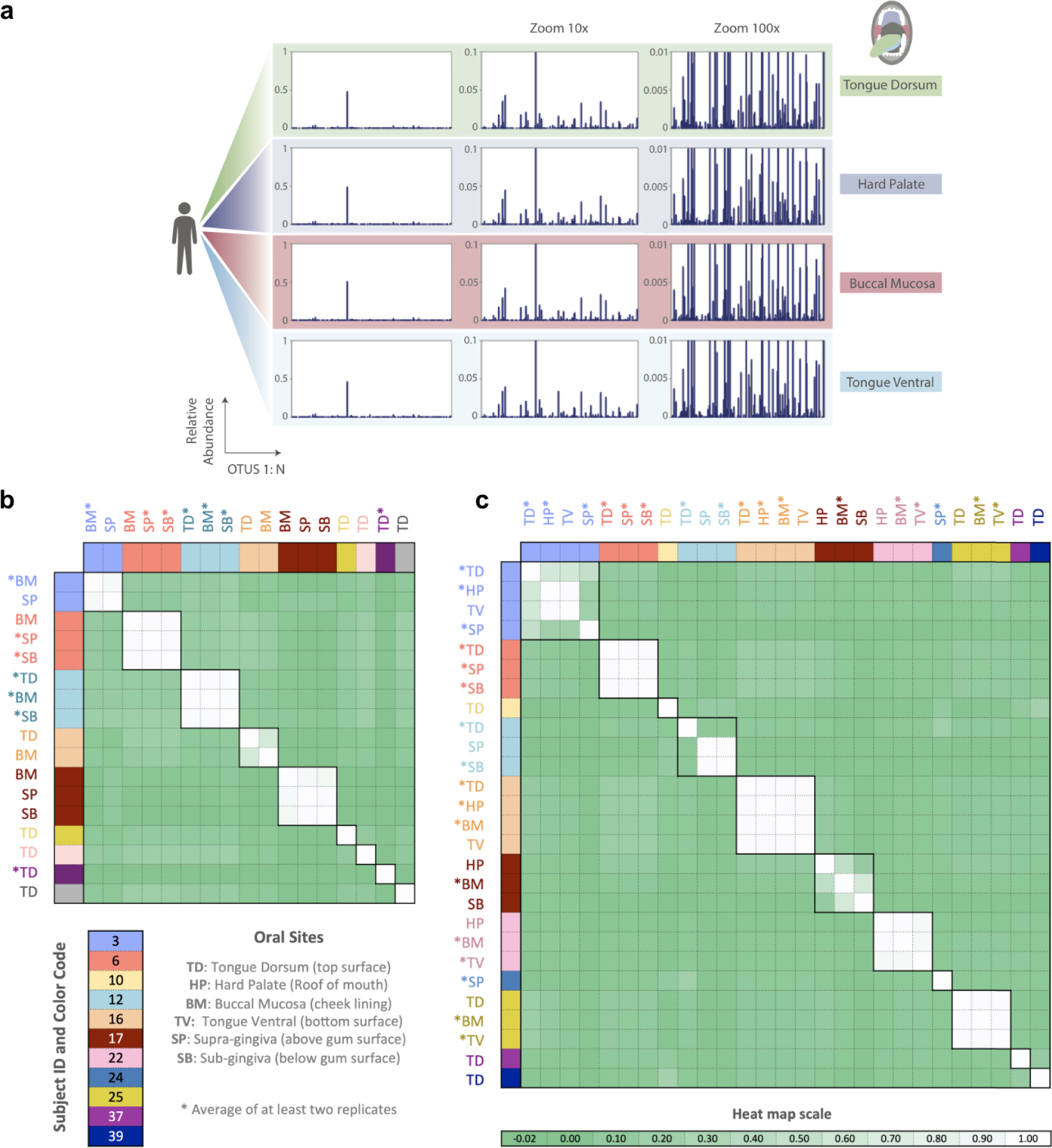
Phageprints across different oral sites. a) HB1 phageprints across four different oral sites in subject 16 zoomed in at 1x, 10x and 100x. b-c) Pearson correlation coefficient matrix of b) HB1 and c) HA phageprints spanning 9 and 11 subjects, respectively. Phageprints are color-coded based on the individual they originate from. Phageprints that have been replicated experimentally at least twice and averaged are denoted by an asterisk. Each phageprint is derived from the analysis of 4000 sequences associated with an individual and a particular oral site. Samples are color-coded based on the individual they originate from. OTUs were defined at 98% sequence similarity and OTUs with less than or equal to 0.001 relative abundance across all phageprints were filtered out. Oral sites shown are the tongue dorsum (TD), buccal mucosa (BM), supra-gingiva (SP), sub-gingiva (SB), hard palate (HP), and ventral surface of the tongue (TV).

As in the case of the HB1 phage family, there is low to non-existing correlation between the HA phageprints of different individuals at the same oral site (**Figure 6**), reinforcing the notion of highly personal phageprints. However, unlike HB1, not all oral sites within the same subject are highly or even moderately correlated (see subjects 3, 12, and 17). In subject 12, for example, the tongue dorsum has a correlation close to zero with supragingival and subgingival sites, which are nearly perfectly correlated. Similarly, in subject 3, the hard palate and the tongue ventral surface have nearly identical phageprints while they have a very low correlation with the phageprint from the tongue dorsum. However, unlike subject 12, the tongue dorsum in subject 3 seems to be an intermediate site, having a moderate correlation with all other sites that are distinct from each other. In subject 17 as well, buccal mucosa serves as the intermediate environment with a phageprint that exhibits a moderate correlation with the disparate phageprints of subgingival and the hard palate. Phage-host network representations for HB1 (**SI Figure 20**) and HA (**SI Figure 21**) phage terminase families across this cohort demonstrates the cause of weak or strong correlations between different oral sites.

## Discussion

This study provides a high-resolution window into two families of phage terminases in the human mouth. It contains a large global cohort of related and unrelated individuals, completely separate from the cohort we used to identify the two phage families through publicly available datasets. We combined the advantages of metagenomics with targeted sequencing to characterize phage terminase families with a resolution that is unavailable through metagenomic studies. As a result, we were able to observe the phage mutant spectra which despite the temporal fluctuations of each variant were largely personal and persistent in time. Cohabitating siblings and even identical twins did not have phageprints that were any more similar than those of unrelated individuals. The only factor we observed that contributes to phageprint relatedness is direct contact between two habitats, as is demonstrated by the similarity between oral phageprints of partners that even have certain instances of concordantly fluctuating variants.

While others have studied the mutant spectra of RNA pathogenic viruses such as SARS-COV2 and RNA phages^31,52^, the mutant spectra of DNA phages, especially *in vivo*, have not been deeply explored. Even though DNA phages have orders of magnitude lower mutation rates compared to their RNA counterparts, we showed that emergence of phageprints is directly the result of the remarkable sequence diversity within DNA phages.

Moreover, previous metagenomic studies have shown that certain features of the human microbiome are stably associated with individuals over extended periods and could be individual-specific^53–55^. One study that has rigorously tested the identifiability of individuals based on bacterial metagenomic datasets (acquired through the Human Microbiome Project), developed “metagenomic codes” based on the principle of a hitting set^56^. As the authors point out, these metagenomic codes are capable of identifying on the order of 100s of individuals. However, they note that instances of false positives rise as more individuals are added^56^. Additionally, metagenomic codes were only able to identify ∼30% of individuals at a second sampling time point taken ∼30-300 days later^56^.

As far as we know, targeted studies that explore the oral phageome and its potential in human identification are missing from the literature. Using a phage-based targeted approach for human identification has a significant advantage. Namely, the space of possible unique barcodes has not been exhausted. We have only sampled a small region of one terminase family using only thousands of sequences per sample. Deeper sequencing of that one region, or the addition of more terminase families, or other phage gene families-of which there are many in the human phageome-can likely generate an astronomical number of unique phageprints from which individuals could potentially be identified.

To provide more concrete estimates, in our study of more than 100 individuals, we found only one case of unrelated individuals whose phageprints had similar correlation coefficients. This is due to the coarse-graining associated with Pearson correlation matrices, where the most abundant OTU significantly impacts the correlation value between two phageprints. As we showed by the removal of the top ten most abundant OTUs and further random downsampling to 2% of the remaining OTUs, even significantly reduced phageprints remain highly predictive of their subject of origin. However, if we conservatively assume that each terminase family can provide just 50 unique patterns or phageprints using one of the coarsest analysis metrics, namely the Pearson correlation, then the combination of phageprints from six short regions (∼300 bp each) of non-homologous terminase families, equivalent to ∼2 Kb of phage DNA, could provide a greater number of possible patterns than the size of the current human population (50^6^, or 15 billion phageprints). Much larger cohorts would be needed to validate these estimates and they are provided as a crude attempt to illustrate the immense diversity of sequences we carry that is not our own.

Moreover, our previously described method for finding ubiquitous human oral phages relied on relatively small metagenomic datasets, which contained sequences from just seven individuals^35^. Yet, on the basis of markers designed from this small dataset we were able to identify the same phage terminase families in at least 10 times as many individuals from across the globe within the HMP cohort as well as this study’s separate global cohort of ∼100 individuals. This finding further confirms that certain phage families are a stable feature of the human oral microbiome. Studies of phages from various natural environments also report the finding of phage families that are distributed across similar types of habitats despite vast geographical distances and barriers that exist between these habitats^57,58^. The ubiquitous presence of the identified phage families in individuals, together with their temporal persistence, seems to suggest that they likely play important roles in this environment.

An important limitation of metagenomic based discovery of phage families is that host information is often deduced via homology searches between phage sequences and prophages within publicly available bacterial genomes. Gaining more exact information about the host dynamics is very challenging because it first requires the development and validation of species-level or perhaps strain-level bacterial markers - as opposed to generic makers like 16S-that would exclusively target the host species of the HB1 and HA phage families. Even with such markers at hand, untangling the specific host of a particular phage variant without single-cell isolation will be difficult at best. This is because, within a single infection event, one could expect numerous variants and so there will not be a one-to-one pairing of phage and bacterial variants. Single-cell single-phage studies will likely provide the necessary platform for gaining precise pairing of phage and host variants, however these studies do not lend themselves to the spatio-temporal exploration of phages and their hosts in their native contexts. Thus, new methodologies are needed to provide high-resolution exploration of bacterial and phage dynamics without the loss of contextual and population-level information.

## Conclusion

Metagenomic studies have transformed our understanding of phages, though they often fall short in providing the depth of sequencing needed to observe the population structure and dynamics of individual variants within a phage family. Through targeted sequencing of two non-homologous oral phage terminase families, we show that the abundance profile of variants within each family provides unique individual-specific and temporally persistent barcodes or “phageprints”. Analyzing oral samples from ∼100 individuals across several continents, we observed consistent trends: dominant variants coexist with less abundant ones. Machine learning enabled precise differentiation of individuals in the temporal cohort, even when phageprints were downsampled to contain only a small subset of the more rare variants. Notably, factors like residence, genetics, and diet do not appear to impact phageprint similarity. We showed that minimal spatial differences within oral sites produced site-specific phageprints for one of the terminase families. Moreover, through daily monitoring of phageprints, we identified the highly dynamic yet persistent variants and showed that partners’ shared variants can have concordant temporal fluctuations.

## Materials and Methods

### Subject recruitment and sample collection

For the bulk of our sample collection, we relied heavily on citizen scientists. We made an educational video to introduce a diverse audience to the fascinating world of phages, explain our study and to recruit volunteers. We also created an instructional video for prospective volunteers on subject disqualifying criteria and subject rights, and to provide a step-by-step demonstration of sample collection, storage, and shipment. Among other exclusion criteria, subjects could not have taken antibiotics for the preceding 3 months and subjects could not have active cavities or gum disease.

Qualified subjects were sent a kit and were asked not to brush their teeth or tongue for a minimum of 8 hours prior to sample collection to allow for a substantial build up of plaque on the tongue dorsum. Subjects were instructed to 1) wear gloves, 2) scrape their tongue (dorsal surface) several times using the tongue scraper, 3) deposit their sample into the collection tube, 4) place the tube back into the bag, and 5) store the bag in their freezer along with ice gel packs prior to an overnight shipment of their samples. They were also instructed to report any sources of error that occurred at any step, and to send their samples along with their signed consent form and questionnaire. For the temporal study, subjects failed to sample all 30 days. The mean and median number of sampling days were 25. Our sample collection and processing protocols were approved by Caltech Institutional Review Board (IRB protocol 14-0430) and Institutional Biosafety Committee (IBC protocol 13-198).

Nine subjects included in this study are those included in a previous study of oral microbial diversity by Bik *et al*. (21). Briefly, samples were collected from individuals by a dentist who examined subjects for their oral health, thereby excluding subjects with active cavities, gingivitis, or periodontal disease. For each subject, samples from different oral sites were collected using sterile curettes and deposited separately in 1.5 mL collection tubes containing PBS buffer. The 6 oral sites sampled include plaque from tongue dorsum, tongue ventral, buccal mucosa, hard palate, supra-gingiva, and sub-gingiva.

### The sample collection kit and measures against sampling contamination

To obtain samples, we developed a sample collection kit and prepared kit contents within the PCR flowhood. Before and after every kit preparation session, the flowhood surfaces and pipettes were wiped using sterile wipes, DNA AWAY™, and 95% ethanol. At the end of each session the surfaces were also UV-sterilized (60 minutes). Each kit contains plastic tongue scrapers (Yellow CeraSpoon Safe Ear Curettes, Bionix) that were first autoclaved and then UV-sterilized for 60 minutes, 1.5 mL gamma-sterilized and pre-packaged collection tubes certified as pyrogen-RNase-DNA- and ATP-free (VWR), each containing 200 uL sterile 1X PBS buffer (VWR), along with pre-packaged sterile gloves (VWR). Each collection tube and tongue scraper pair was placed inside a sterile bag and the bags were placed in another bag. The next steps were performed outside of the flowhood. Each collection bag was put inside a Styrofoam box along with ice gel packs. Ice gel packs and Styrofoam boxes were not reused to prevent cross contamination between individuals in case of a spill, which would already be highly unlikely due to multiple layers of packaging. Upon arrival of samples, collection tubes were taken out of their original bags, wiped with 95% ethanol and DNA AWAY™ using sterile wipes and placed into a new sterile bag. Gloves were frequently exchanged both during this step and before proceeding to the next collection tube to prevent cross contamination. In addition to standard lab attire such as gloves and lab coat, a face mask was worn to prevent contamination during kit preparation and sample storage.

### Measures against PCR and DNA extraction contamination

A common source of contamination in PCR originates from previously amplified template sequences that enter new PCR reactions. To prevent this type of contamination, four physically separated workstations were developed for DNA extraction (station A1), PCR preparation (station A2), PCR and gel electrophoresis (station B1), and PCR cleanup (station B2). A and B specify two different buildings at Caltech while 1 and 2 refer to two different rooms within the same building. The flow of materials was from building A to B and never vice-versa. Every station had its own set of lab equipment, materials, and storage space. Disposable lab coats (Sigma-Aldrich®) were worn and disposed of at the end of every procedure to ensure that DNA was not carried between stations via clothing. Facemasks (Fisher Scientific) were also worn at all times to prevent any oral or nasal droplets from entering reactions. Prior to the start of every DNA extraction, lab equipment and bench tops were cleaned using sterile wipes and DNA AWAY™ (Thermo Scientific), a surface decontaminant that eliminates DNA and DNAses. PCR preparations and aliquoting of reagents were carried out in a PCR flowhood (AirClean® Systems) equipped with a UV light and laminar airflow capabilities. Lab equipment required for PCR preparation was designated to the PCR preparation flowhood. At the end of every experimental session and when introducing new equipment into the flowhood, all surfaces were first wiped with DNA AWAY™ solution and then exposed to UV radiation for 60 minutes. Prepackaged, sterile gloves were used for PCR preparation. To prevent sample-to-sample contamination during DNA extraction, PCR preparation, and PCR cleanup, gloves were frequently exchanged. Most importantly, 5 No Template Control (NTC) reactions accompanied every PCR run. Similarly, to test the presence of contaminants in extraction reagents, for every extraction experiment, 3 reactions were carried out without the addition of any sample. PCR using phage primers was performed on these extraction control reactions.

### DNA Extraction (Station A1)

DNA extraction of human oral samples was done according to the manual from MoBio PowerBiofilm® DNA Isolation Kit. The advantage to using this kit for DNA extraction and purification is that it combines the use of chemical and mechanical (bead-beating) treatments for an increased efficiency in biofilm disruption, lysis, and removal of inhibitors such as humic acid. The final concentrations of DNA were measured using Nanodrop. The concentration range of the total extracted genomic DNA was typically between 5 to 50 ng/µL.

### PCR preparation (Station A2) and PCR (Station B1)

Each PCR reaction contained 12.5 µL of PerfeCTa® qPCR SuperMix, ROX™ (Quanta Biosciences), a premix containing AccuStart™ Taq DNA polymerase, MgCl_2_, dNTPs, and ROX reference dye for qPCR applications. Additionally, each reaction contained 10.5 µL of RT-PCR Grade Water (Ambion®) which is free of nucleic acids and nucleases, 1 µL of extracted DNA at 1 ng/µL, 0.5 µL of forward and 0.5 µL of reverse primers, each at 50 ng/µL (synthesized by IDT). A higher than recommended primer concentration was used because the phage primers used are 32-64 fold degenerate. The thermocycling protocol was made according to PerfeCTa qPCR SuperMix recommendations: 1) a 10-minute activation of AccuStart™ Taq DNA polymerase at 95°C, 2) 10 seconds of DNA denaturation at 95°C, 3) 20 seconds of annealing at 60°C, and 4) 30 seconds of extension at 72°C, 40 cycles repeating steps 2 to 4, followed by 5 minutes of final extension at 72°C.

### Gel electrophoresis (Station B1) and PCR cleanup (Station B2)

Phage PCR products were visualized using 2% agarose in TAE buffer. After gels were cast, 5 µL of each PCR product was mixed with 1 µL of 6X loading dye and loaded into a well. 5 µL of 100 base-pair ladder was used, and the gel electrophoresis instrument was set to run for 30 minutes at 100V. Phage PCR positive hits were purified using the QIAquick PCR Purification Kit (QIAGEN). 20µL of PCR products were used and purified according to the QIAquick PCR Purification manual.

### Sequencing

Upon PCR cleanup, double stranded DNA concentration in each sample was measured using the Qubit instrument. Qubit measurements were performed in Building C due to practical considerations rather than a necessary treatment for preventing contamination. Samples were combined into one reaction (∼2 μg dsDNA) and submitted to GENEWIZ, Inc for library preparation and MiSeq 2×300bp Paired-End sequencing. To enable multiplexing, unique DNA barcodes (Table 1) were appended onto the forward primer sequences (Table 3) used to amplify each phage terminase family. These barcoded primer sequences were synthesized by IDT. Using this scheme, ∼100 samples were submitted per MiSeq sequencing run (Table 1) and by matching the barcode sequence to the sample ID, information about who and where the sample came from was accessible. More specifically, Hamady error-correcting 8-letter barcodes^59^ were used. Hamady DNA barcodes are an example of Hamming code wherein the addition of parity bits allow for detection and correction of errors within the barcode sequence. In the case of Hamady barcodes, up to 2 errors in the barcode sequence can be detected and one error can be corrected.

### Quality control steps to eliminate sequencing errors

We used Illumina MiSeq’s 2×300bp paired-end configuration (GENEWIZ, Inc). Each sequencing run produced about 20-25 Million paired-end reads. Paired-end reads were joined using *join_paired_ends.py* script from QIIME (Quantitative Insights Into Microbial Ecology)^60^ package, and unless noted otherwise scripts used are part of QIIME. If paired reads had any mismatches across their overlapping bases, the paired reads was eliminated from any further analysis (QC step #1). The overlap between the paired reads nearly entirely covers the HA and HB1 sequences (∼300 bases long), hence eliminating most sequencing errors.

Upon joining reads and eliminating those with mismatches in the region of overlap, *seqQualityFilters.py*, an in-house script, was used to perform QC step #2: taking joined reads from QC step #1, and eliminating any sequences that have one or more bases marked by a Phred score below 30. Excluded from QC step #2 were the first two bases in the beginning and end of each sequence, which for the majority of reads have much lower quality scores.

Using *seqQualityFilters.py,* sequences were placed in 2 different bins according to their primer sequences, and any sequence that did not have the correct barcode length, or the correct primer sequences at the expected positions, was eliminated (QC step #3). Additionally, nearly half of remaining sequences had to be reverse complemented so that all sequences were oriented in the 5’ to 3’ direction. Using the same script, primer and barcode sequences were removed, and barcode sequences were written to a separate file (to be used as input to *split_libraries_fastq.py*). At this point sequences that did not have the correct length were filtered out (QC step #3). Sequences were demultiplexed using *split_libraries_fastq.py* and reads with errors in the barcode sequence were eliminated (QC step #4).

### Phage terminase OTU relative abundance plots or “Phageprints”

After demultiplexing quality-controlled reads, sequences were clustered according to a specified sequence similarity threshold using UCLUST *de novo* clustering algorithm^61^ used in *pick_otus.py* script. Using *make_otu_table.py*, OTU tables were generated. An OTU table summarizes counts of sequences assigned to each OTU across each sample. We refer to this per-sample sequence count as the OTU size. As long as an OTU of size 1 or greater exists in at least one sample, it is included in the OTU table. In this way, the counts of OTUs for samples containing the same phage terminase family remains the same, though their size could vary widely across different samples. The relative abundance of each OTU within each sample was calculated via *processOtuTable.py,* another in-house script. In plotting the relative OTU abundance values for different samples, we arrived at complex, individual-specific patterns. We named these plots “phageprints”.

The most abundant sequence from each OTU was retrieved using *pick_rep_set.py* to serve as a representative sequence. HB1 representative sequences were aligned using Geneious, using a gap open penalty of 30 and gap extension penalty of 15 and a 65% similarity cost matrix. No gaps were introduced. The alignment is shown in **SI Figure 22**.

### Examining the effect of OTU sequence similarity threshold

In analyzing 16S sequences, clusters or Operational Taxonomic Units (OTUs) are conventionally defined at 97% sequence similarity threshold. To examine the effect of sequence similarity threshold for phage OTU formation, we tested OTU sequence similarity thresholds of 98%, 97%, 95%, 90%, and 80%. **SI Figure 3** is a matrix of Pearson correlation coefficients calculated during the pairwise comparison of HB1 phageprints using different sequence similarity thresholds for defining OTUs. Very similar Pearson correlation matrices are obtained as the sequence similarity threshold is lowered from 98% to 80%. However, because the number of clusters is reduced as we reduce the sequence similarity threshold, with lower sequence similarity thresholds, the chance that individual-specific variations are lumped into the same cluster is increased. If noise-induced sequence variations are effectively accounted for, higher sequence similarity thresholds for defining OTUs can enable a more accurate and detailed depiction of a person’s phageprint. For this reason, we used a sequence similarity thresholds of 98% or 100% in this study.

### Detecting experimental noise

How reproducible is a phageprint plot? **SI Figure 4** summarizes the sources of noise from all experimental processes performed during this study. First, it’s important to capture sampling variation. How consistently can we capture a phageprint from an individual’s oral site given that we are sampling different parts of the biofilm each time? Another factor that could contribute to sampling variation are the personal differences in the rate of biofilm mass accumulation on the tongue dorsum. Secondly, we need to ask whether processes of lysis and DNA extraction allow for the availability of the same template DNA sequences in the same relative abundances across different extraction runs.

Moreover, we need to evaluate the OTU abundance variations that could result in PCR due to both errors as well as other stochastic events. For example, it’s possible that very rare template sequences are left out of the initial cycles of PCR and their relative abundance at the end of PCR is lower than their relative abundance prior to PCR, and thus PCR could serve as a biased amplifier. PCR purification is similar to extraction and sampling in that it does not introduce sequence errors; however it is unlike these processes because after PCR billions of template copies are created and it’s unlikely that the loss of a fraction of templates during PCR purification will dramatically change OTU relative abundances. Finally, Illumina MiSeq sequencing is another error-prone process not only at the level of base-calling, but at the level of bridge amplification.

To quantify how reproducible a given phageprint is, we obtained 3 different samples from a subject’s (subject 37) tongue dorsum. We then performed DNA extraction and PCR separately on each sample and sent samples for sequencing (sequencing run #2). The logic behind this experiment was to capture a lumped measure of noise arising from various processes depicted in **SI Figure 4**. After performing quality control steps 1-4, demultiplexing reads based on their barcode sequences, clustering reads based on 98% sequence similarity threshold for OTU formation, rarefying the OTU table to 4000 reads per sample, and calculating the relative abundances of OTUs, we measured the standard deviation in the relative abundance of each OTU across these three samples (**SI Figure 5**). We show that the relative abundance values across these three samples are highly consistent, with the majority of OTUs having standard deviations below 0.002 and the maximum standard deviation observed was ∼0.007 relative abundance.

### Identifying non-reproducible OTUs and OTU relative abundance detection threshold

To identify OTUs that were non-reproducible across the three samples from subject 37’s tongue dorsum (HB1 marker), we flagged OTUs that had appeared in only one or two samples out of three. We then plotted the histogram of non-reproducible OTUs as a function of their relative abundance (for those OTUs appearing in 2 out 3 samples, the higher relative abundance value was used). The thresholds defining each bin, *b*, were selected to be the following: 0>b_1_≥0.00025 (OTU of size 1 sequence since the total number of sequences per sample is 4000), 0.00025>b_2_≥0.0005 (2 sequences), 0.0005>b_3_≥0.00075 (3 sequences), 0.00075>b_4_≥0.001 (4 sequences), and 0.001<b_5_ (5 or more sequences). We demonstrate that the number of non-reproducible OTUs drops as a function of OTU relative abundance, and all OTUs with more than 4 sequences (0.001 relative abundance) are reproducible (**SI Figure 6**). To conclude, we arrived at 0.001 relative abundance as the detection threshold for OTUs.

In addition to capturing a lumped sum of noise across all experimental processes for subject 37 tongue dorsum sample, for samples from subjects 3, 6, 10, 16, and 17, we performed a second set of PCR on previously extracted DNA samples, and submitted those samples for sequencing (**SI Figure 7**). In addition to these replicates, we acquired new samples from the tongue dorsum for subjects 31, 35, 37, and 38, and submitted these samples for the second sequencing run. Overall, we were able to demonstrate that with the quality filtration steps we developed, phageprints are highly reproducible even when they are generated from different PCR and sequencing steps (**SI Figure 7**).

### Phage-host networks

OTU tables were input to *createNetwork.py,* an in-house script that creates node and edge tables. The nodes represent samples and phage OTUs, and a directed edge connects samples to the OTUs that they host. The weight of this connection is based on the relative abundance of the OTU in that sample. Gephi^62^ was used to visualize the resulting networks, and to obtain the degree distribution.

### Machine learning models

We used the scikit-learn^63^ package within Python to create different machine learning model types and report performance metrics such as AUPR and AUROC. The input to these models were various versions of the HA and HB1 OTU tables, containing samples as rows and OTUs as columns. To obtain 95% confidence intervals of performance metrics for a given model type, we used random splits to create 10 different models based on 10 different instances of training (70% of samples) and testing (30% of samples) portions of the input dataset. No extensive parameter tuning was explored as default parameter values already resulted in very high model performances. We provide our notebook that delves into building machine learning models along with all other types of analyses presented in this paper (outside of those performed using QIIME) in our jupyter notebooks available (see **Data and Code Availability**).

### Data and Code Availability

All custom python scripts used to process and analyze data is provided in our GitHub repository: https://github.com/gitamahm/phageprint/

## Acknowledgement

We are grateful to members of the Phillips Lab and the Boundaries of Life Initiative for helpful discussions. This study was supported by the National Science Foundation (Graduate Research Fellowship; DGE-1144469), the John Templeton Foundation (Boundaries of Life Initiative; 51250), the National Institute of Health (Maximizing Investigator’s Research Award; RFA-GM-17-002), and the National Institute of Health (Exceptional Unconventional Research Enabling Knowledge Acceleration; R01-GM098465).

## SI Figures

**SI Figure 1.**
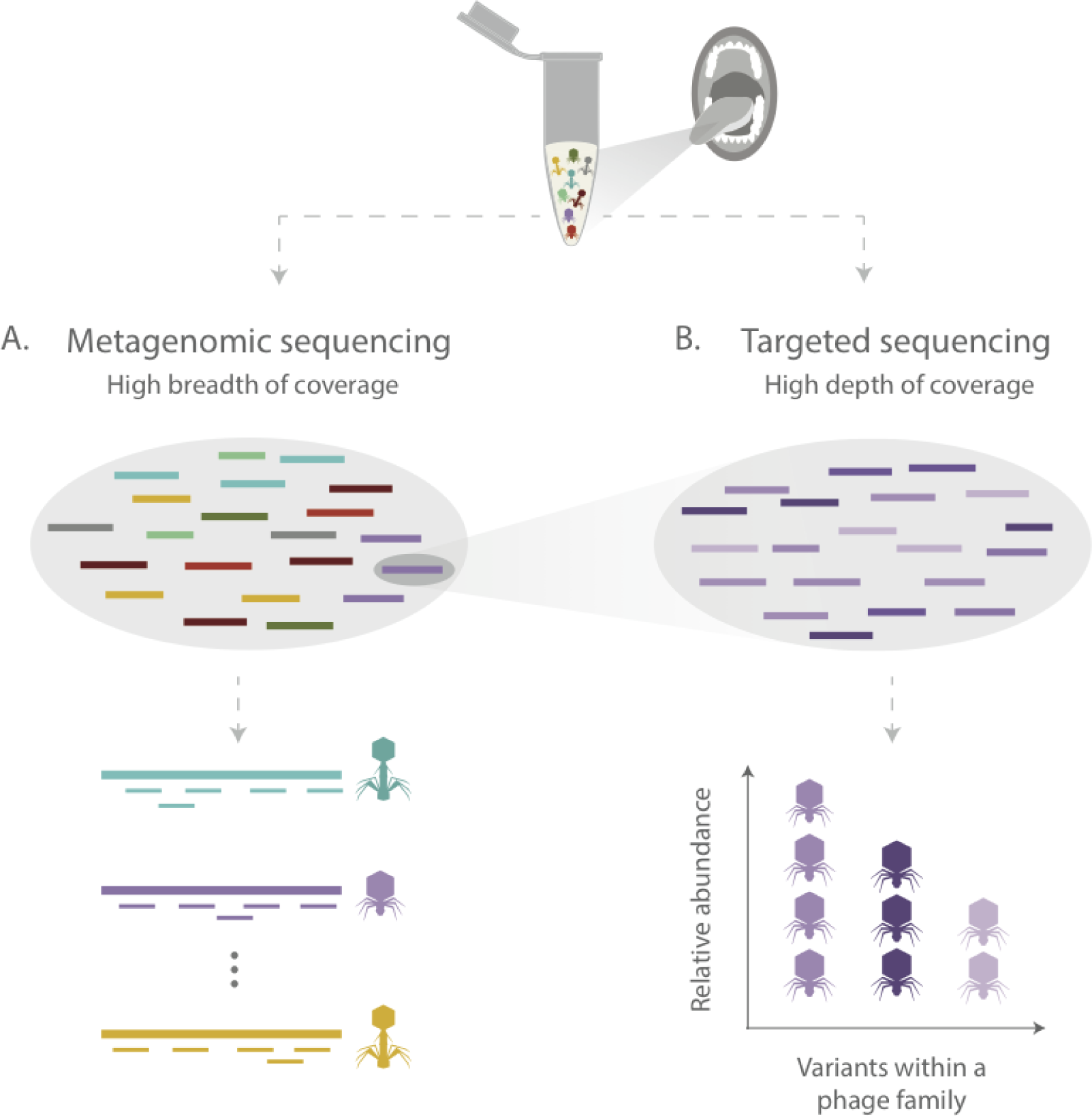
Comparison of A) shotgun metagenomic sequencing and B) targeted sequencing approaches. A) Shotgun metagenomic sequencing offers high breadth of coverage, spanning genomes from many different organisms, however it suffers from low depth of coverage (shown here by the incomplete assembly of phage genomes). B) Targeted sequencing approaches, which use PCR to amplify a specific genomic region, exchange breadth of coverage for depth. Targeted sequencing studies, due to their greater depth of coverage, provide much higher resolution for constructing the communities by equating coverage depth with relative abundance of species or strains.

**SI Figure 2.**
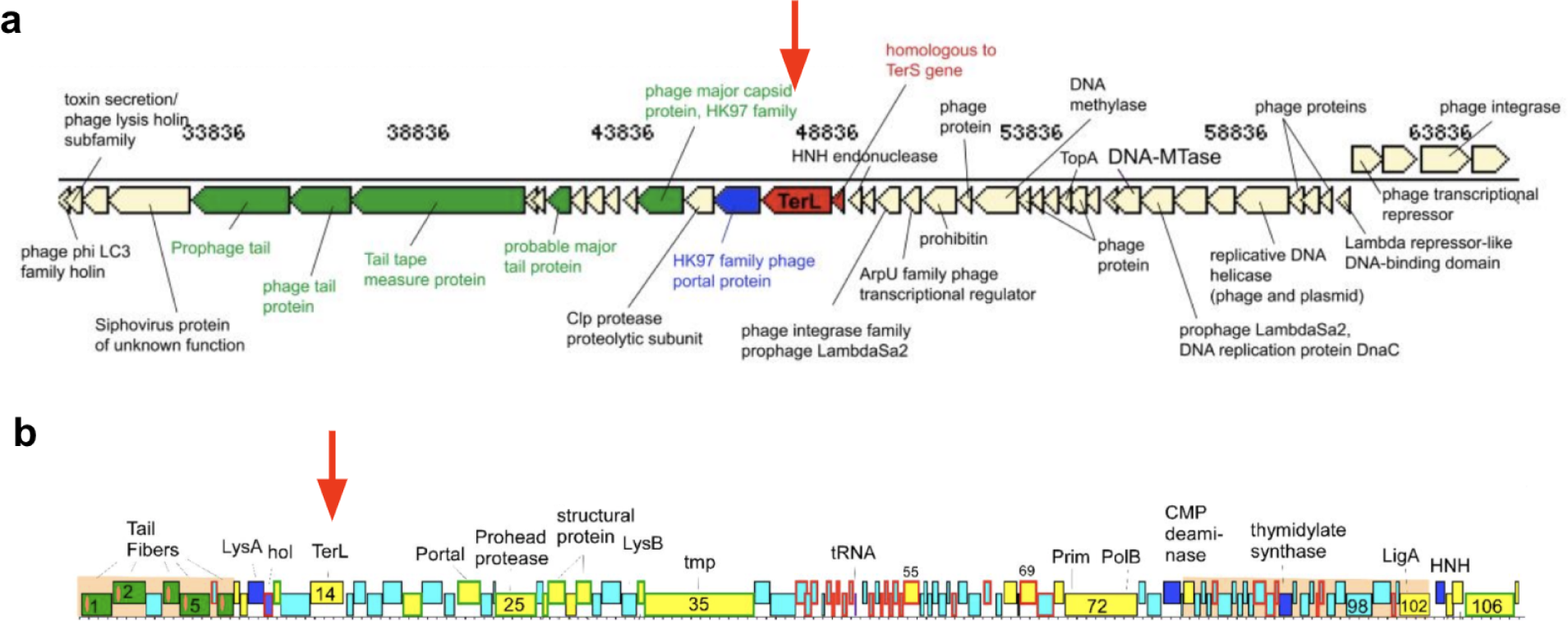
Genomic positions of HA (a) and HB1 (b) homologous terminases found in publicly available databases. a) *Streptococcus oralis ATCC 49296* phage genome containing HA terminase homolog. Figure adopted from our previous study^35^ b) *Rhodococcus equi ReqiPepy6* phage adopted from Summer *et al*^64^. The red arrows point to the position of the large terminase in these phage genomes.

**SI Figure 3.**
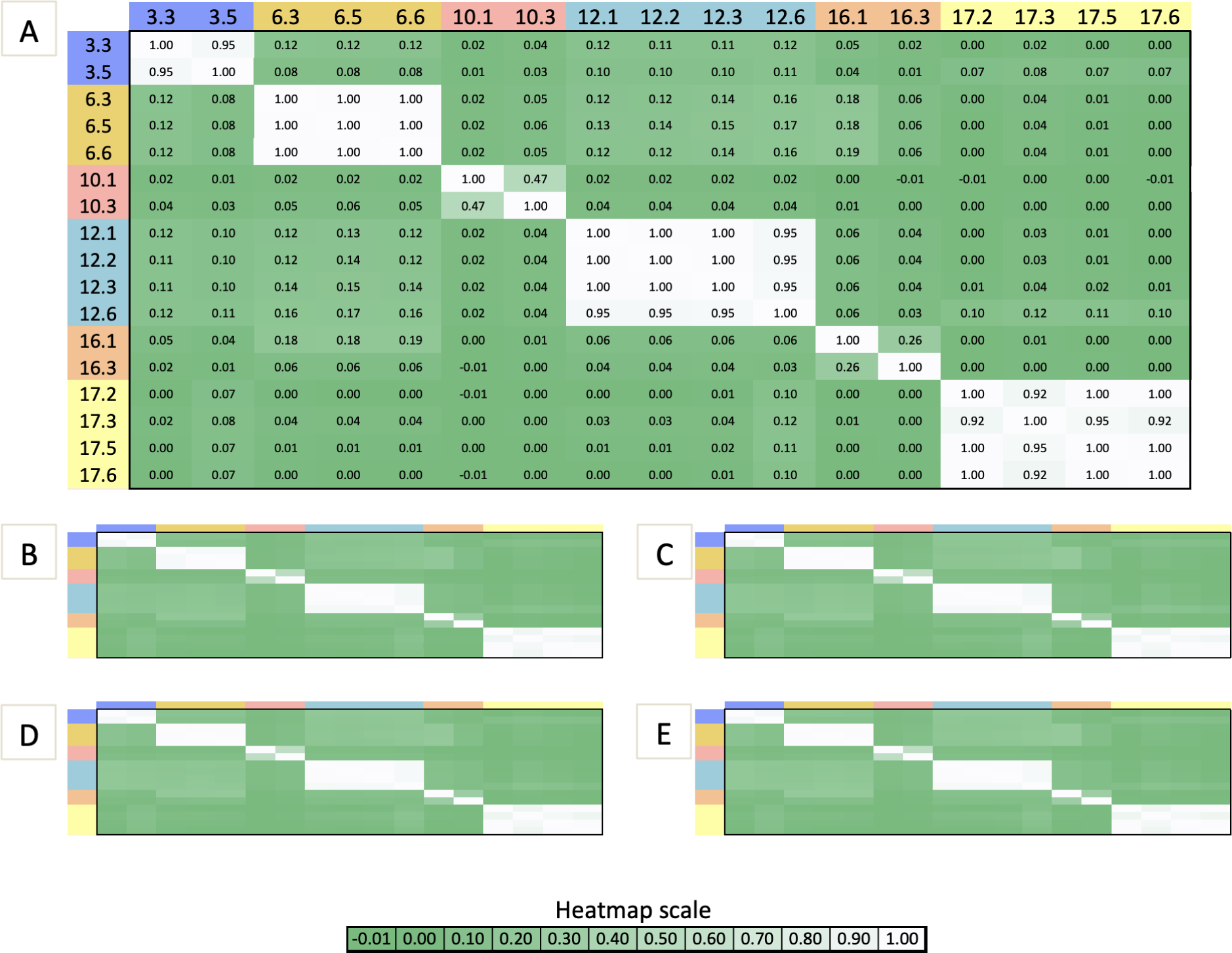
Pairwise Pearson correlation coefficient values calculated for HB1 phageprints as a function of A) 98%, B) 97%, C) 95%, D) 90%, and E) 80% sequence similarity thresholds for OTU formation. Sample IDs can be decoded as such: subject ID precedes oral site ID. Oral sites 1-6 correspond to tongue dorsum, hard palate, buccal mucosa, ventral tongue, supra-gingiva, and sub-gingiva respectively (e.g. 3.3 corresponds to subject 3 phageprint derived from the buccal mucosa, and 3.5 is subject 3 supra-gingiva phageprint). The number of OTUs generated at 98%, 97%, 95%, 90%, and 80% sequence similarity thresholds are 210, 181, 172, 170, and 80, respectively.

**SI Figure 4.**
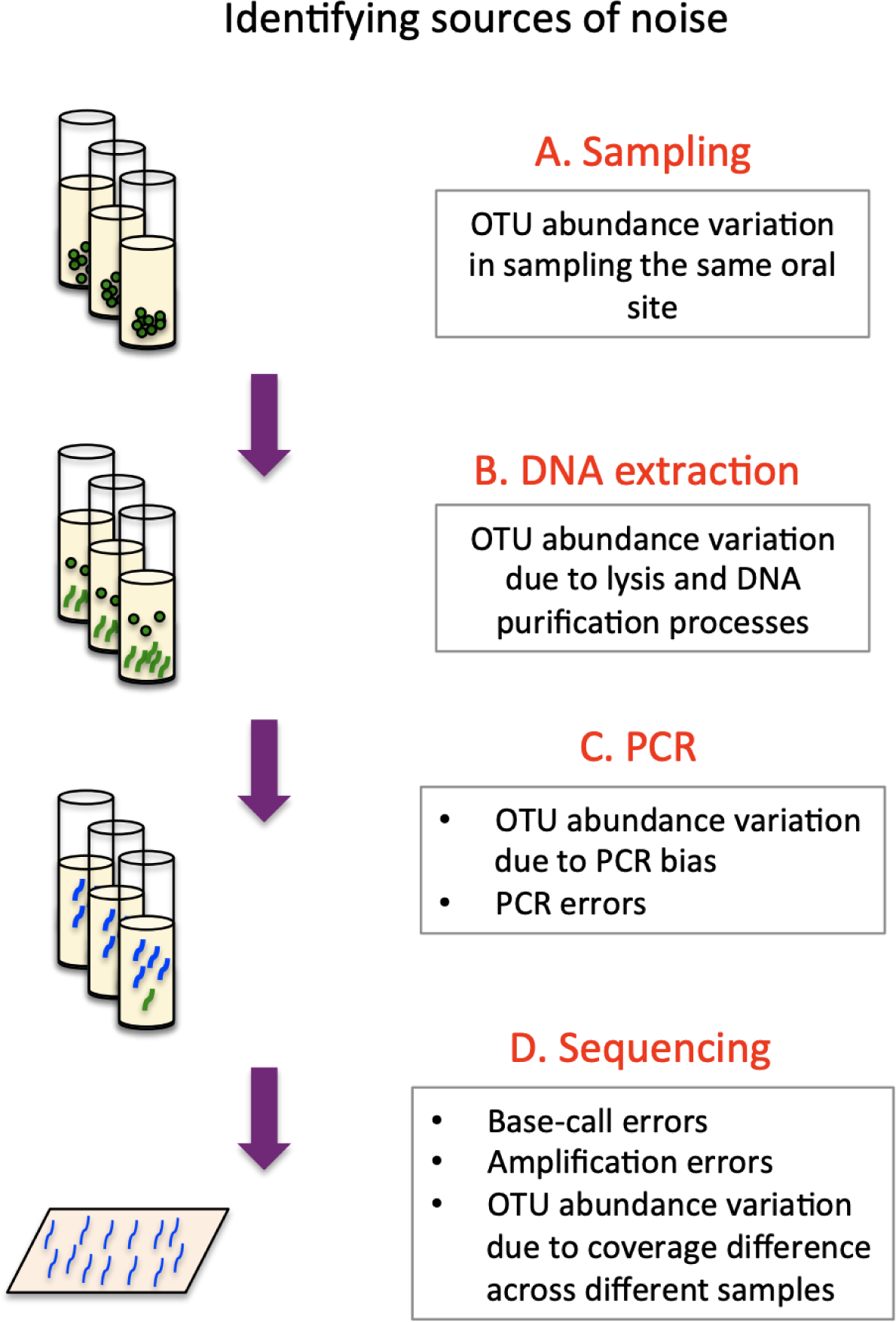
Sources of error and variation in experimental processes used in this study. A) Sampling of the same oral site in the same individual could result in collection of different microbial communities, which could introduce new OTUs or change relative abundance of existing OTUs. B) DNA extraction is not 100% efficient and the fraction of DNA extracted from an environment could serve as a source of variation across different samples. C) PCR introduces errors that could present themselves as novel OTUs or cause variation in abundance of genuine OTUs. D) Sequencing also introduces errors both at the level of base-calling and bridge amplification.

**SI Figure 5.**
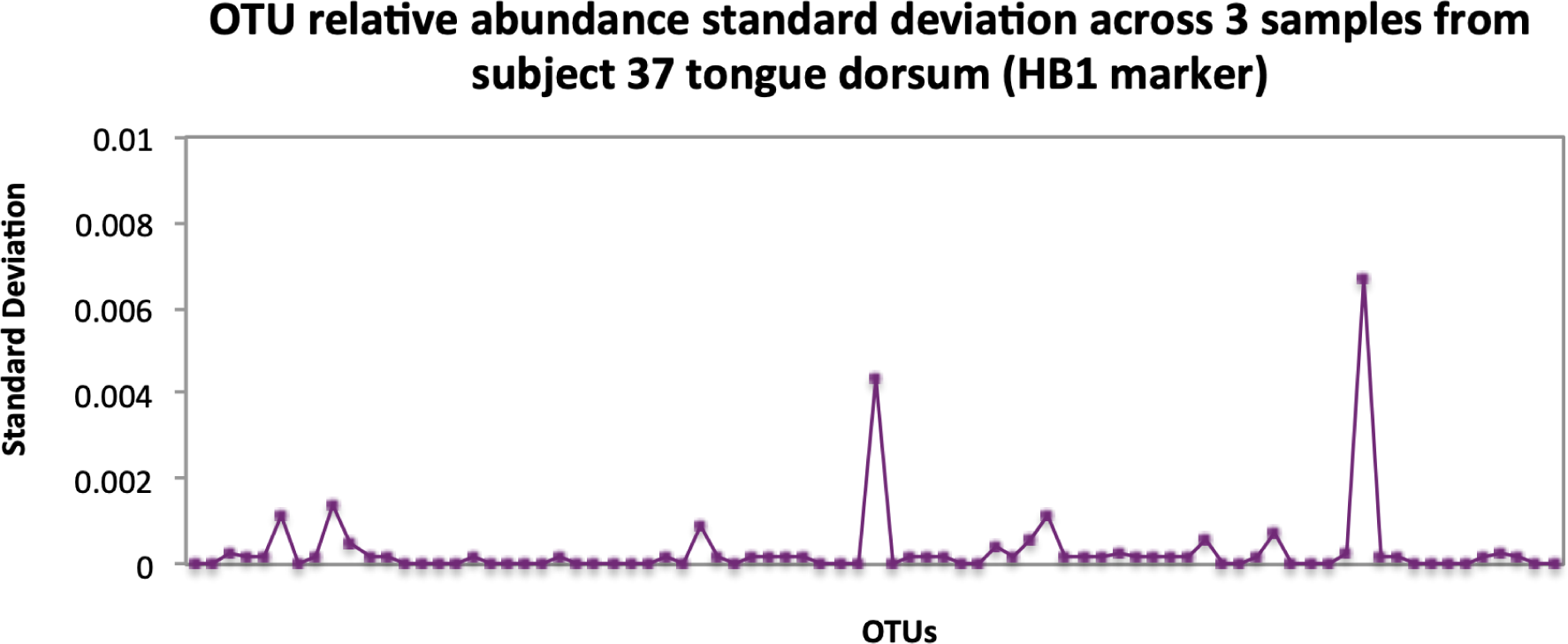
Standard deviations of OTU relative abundances calculated for all experimental processes. Three data points per OTU are used for standard deviation calculations. These three data points correspond to measurements of OTU relative abundances obtained for three different samples obtained from subject 37 tongue dorsum (HB1) which underwent separate sampling, DNA extraction, PCR and PCR cleanup procedures. The maximum standard deviation observed is less than 0.007 relative abundance, and majority are close to 0.

**SI Figure 6.**
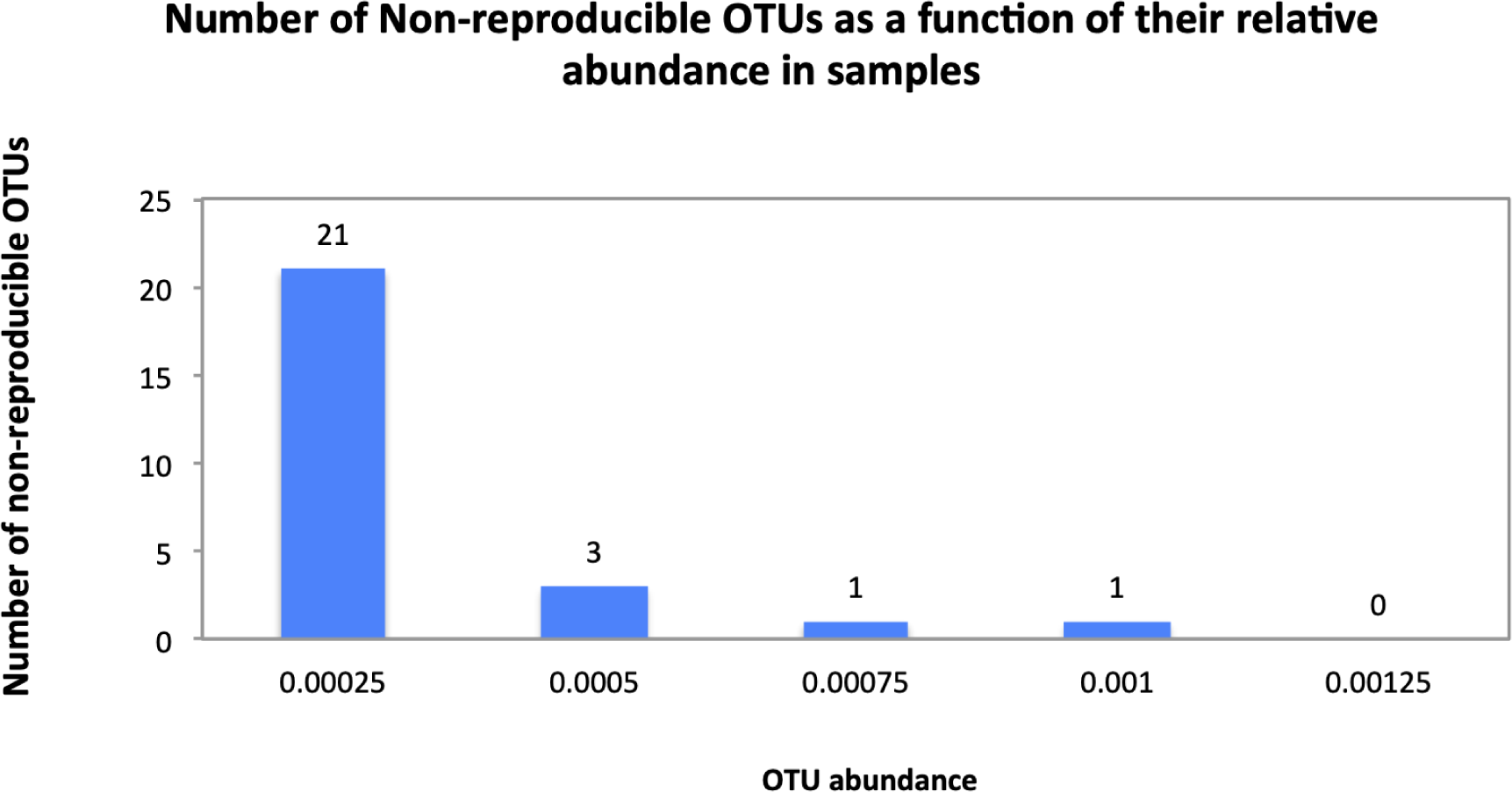
Number of non-reproducible OTUs across three samples obtained from subject 37 tongue dorsum (HB1 terminase family), presented as a function of OTU relative abundance. A total of 30 OTUs appear in one or two samples out of three, and therefore are considered non-reproducible. 21 out of 30 OTUs are defined by a single sequence which translates into 0.00025 relative abundance since samples are rarefied to 4000 sequences. The number of non-reproducible OTUs drops as a function of OTU relative abundance, and all OTUs with more than 4 sequences (0.001 relative abundance) are reproducible across three samples.

**SI Figure 7.**
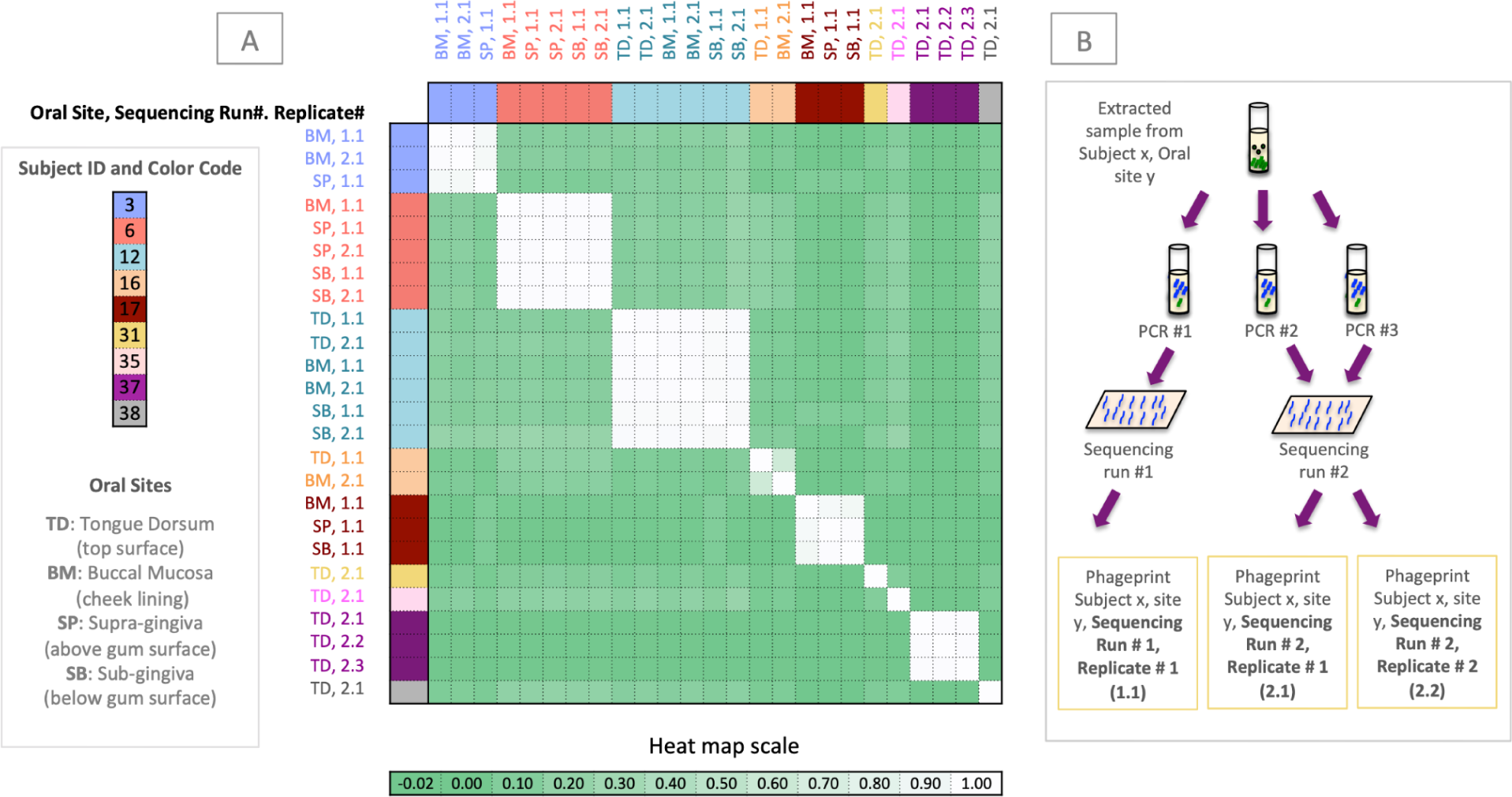
Panel A is the Pearson correlation matrix of all HB1 phageprints. Each phageprint is derived from the analysis of 4000 sequences associated with an individual and a particular oral site. OTUs are defined at 98% sequence similarity and OTUs with less than or equal to 0.001 relative abundance across all phageprints were filtered out. Phageprints are color-coded based on the individual they originate from. Oral sites shown to be positive for the HB1 marker are the tongue dorsum (TD), buccal mucosa (BM), supra-gingiva (SP), and sub-gingiva (SB). Phageprints that were acquired from sequencing run #1, are those marked as replicate #1. Panel B shows that to confirm reproducibility of phageprints, a second set of PCR was performed on previously extracted DNA from all samples included in sequencing run #1 and those PCR products were included in sequencing run #2. Phageprints derived from the second sequencing run are marked as replicate #2.

**SI Figure 8.**
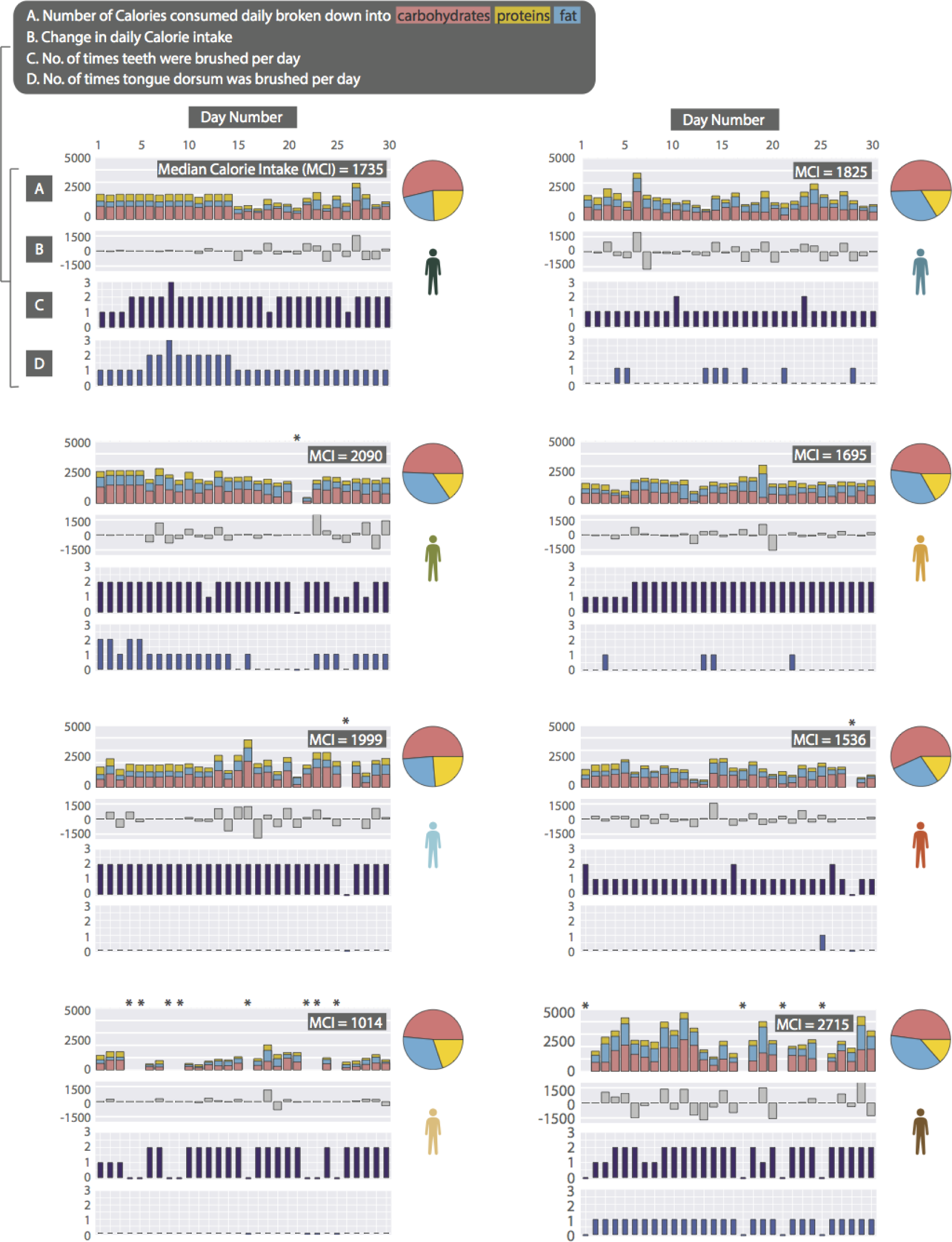
Subject daily metadata for the temporal cohort. Top panel for each subject represents the Caloric intake from fats, carbohydrates, and protein. Mean Caloric Intake or MCI reports the Caloric intake averaged over the sampling period (the x-axis for all plots is number of days). Pie charts demonstrate the diet over sampling days based on median fat, carbohydrate and protein consumption. The second panel depicts the change in Calorie intake from the previous day. The third and fourth panel correspond to the number of times that the subject brushed his or her teeth and tongue, respectively, during the 24 hour sampling interval. We have used an asterisk to denote days for which we did not receive data from the subject, and to distinguish them from zero values in the third and fourth panel, they have been given “-.1” value. Two subjects did not report dietary information so they are not included in this figure.

**SI Figure 9.**
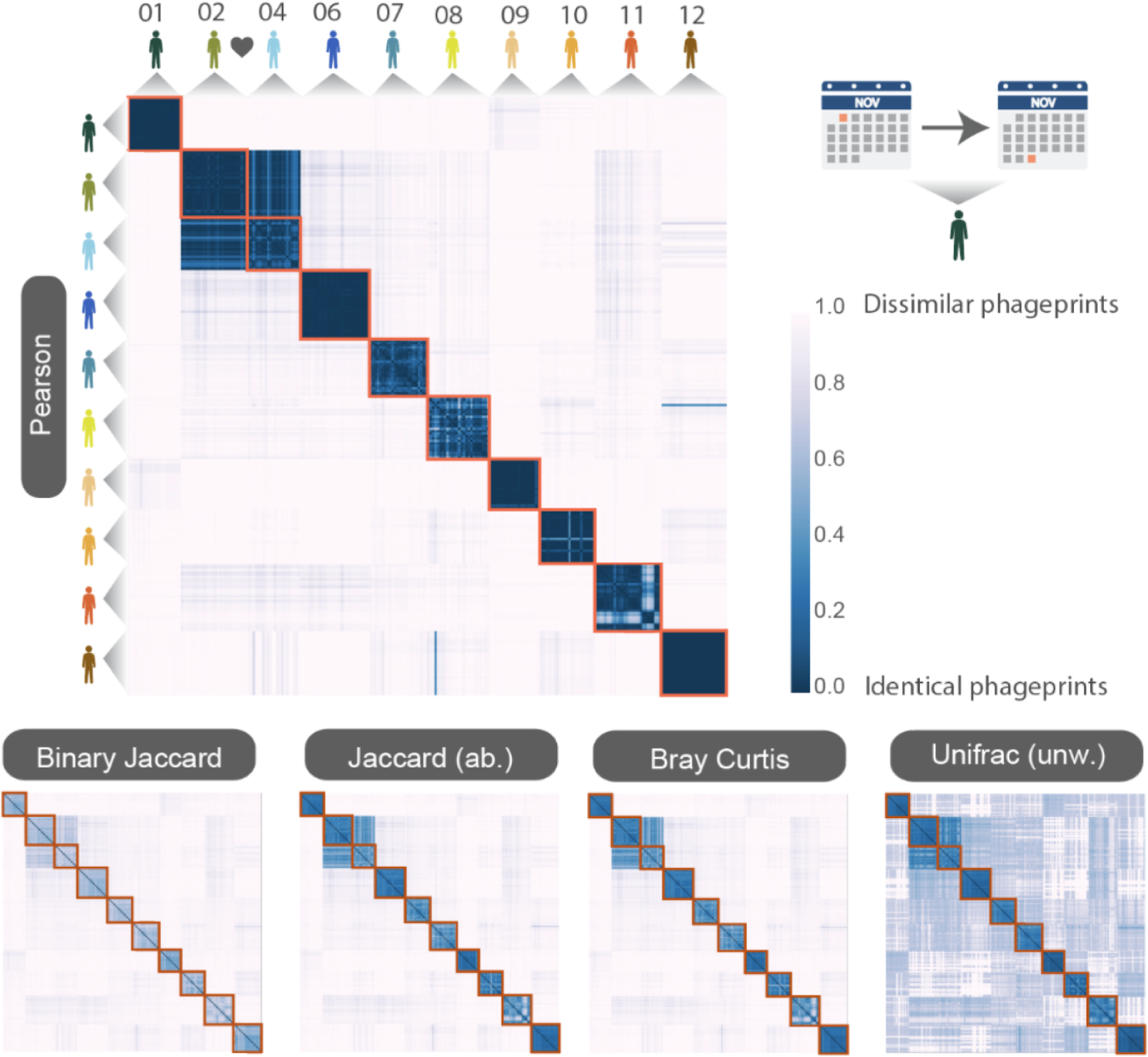
HB1 phageprints temporal dynamics depicted here by pairwise distance metrics: A) Peason, B) Binary Jaccard, Abundance Jaccard, Bray Curtis and unweighted Unifrac. The heatmap scale applies to all heatmaps shown. Subjects 02 and 04 are partners. Samples from each subject are chronologically ordered.

**SI Figure 10.**
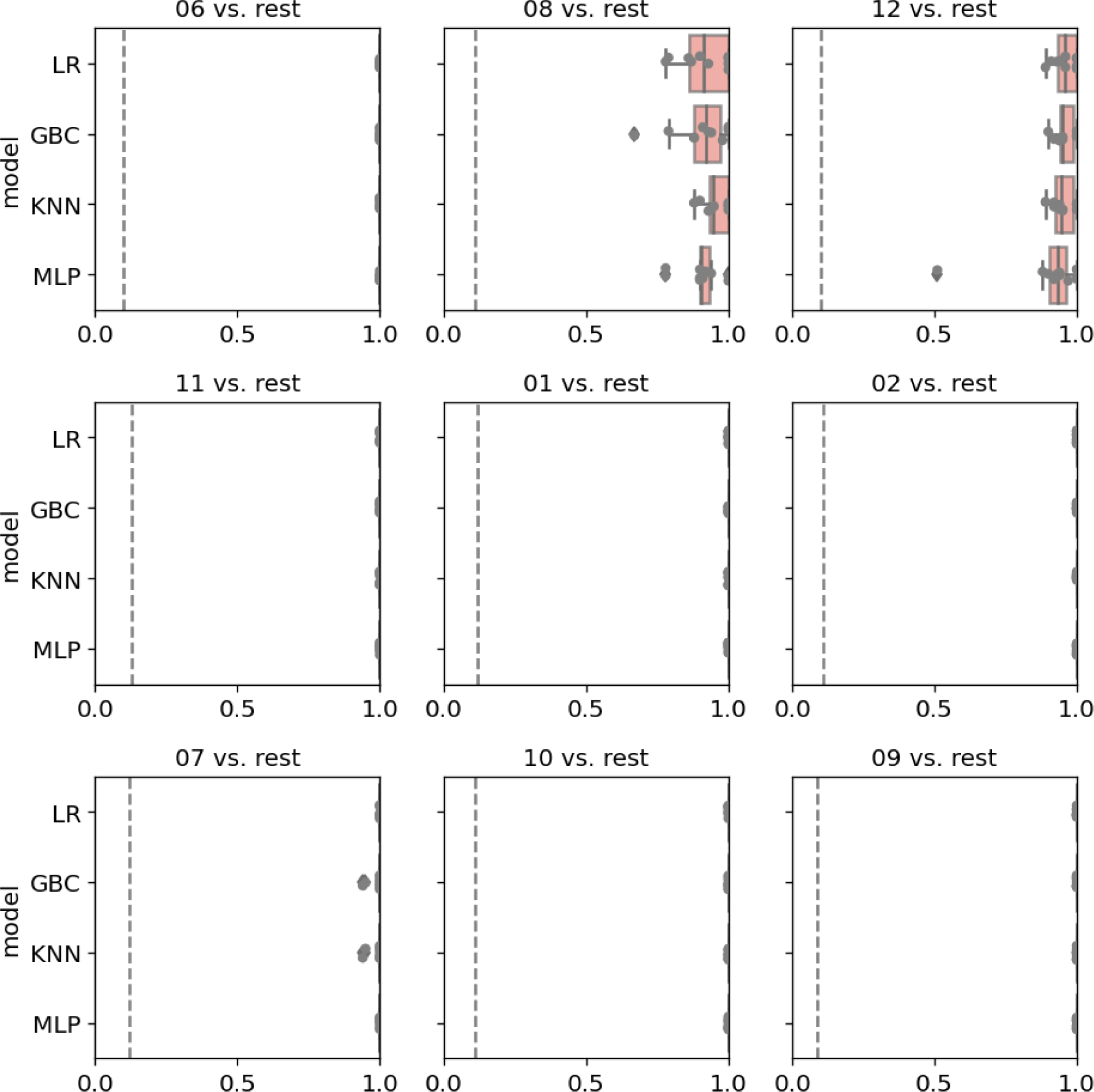
Four types of machine learning models are built to classify one individual’s phageprints from the rest (HA phageprints). For each model type, 10 independent models based on 10 different train/test splits are built, and boxplots of the Area Under the Precision Recall Curve (AUPR) is reported in this table. The null value (i.e. prevalence of the positive class) is shown as a dashed line.

**SI Figure 11.**
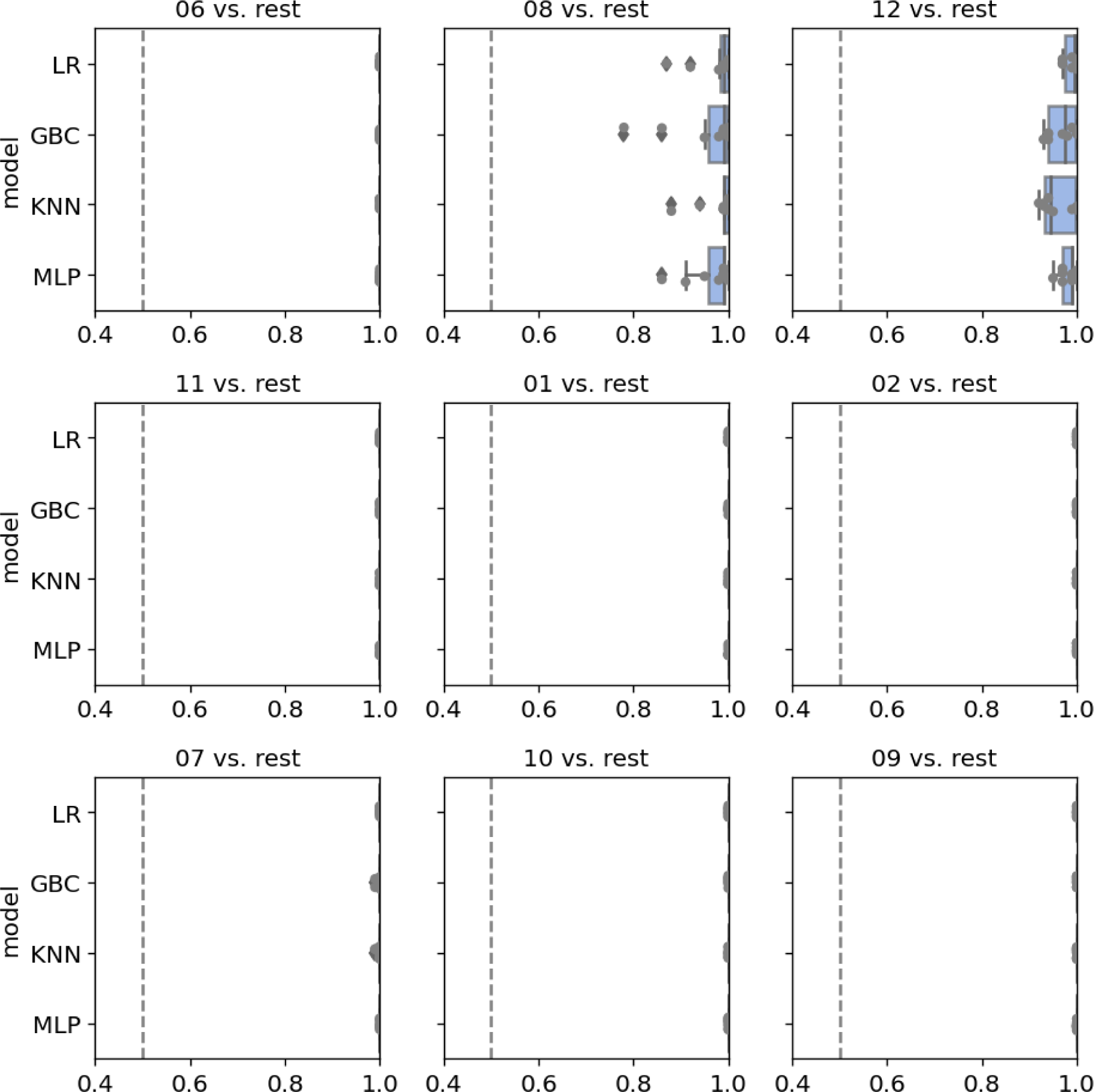
Four types of machine learning models are built to classify one individual’s phageprints from the rest (HA phageprints). For each model type, 10 independent models based on 10 different train/test splits are built, and boxplots of the Area Under the Receiver Operator Curve (AUROC) is reported in this table. The null value (0.5) is shown as a dashed line.

**SI Figure 12.**
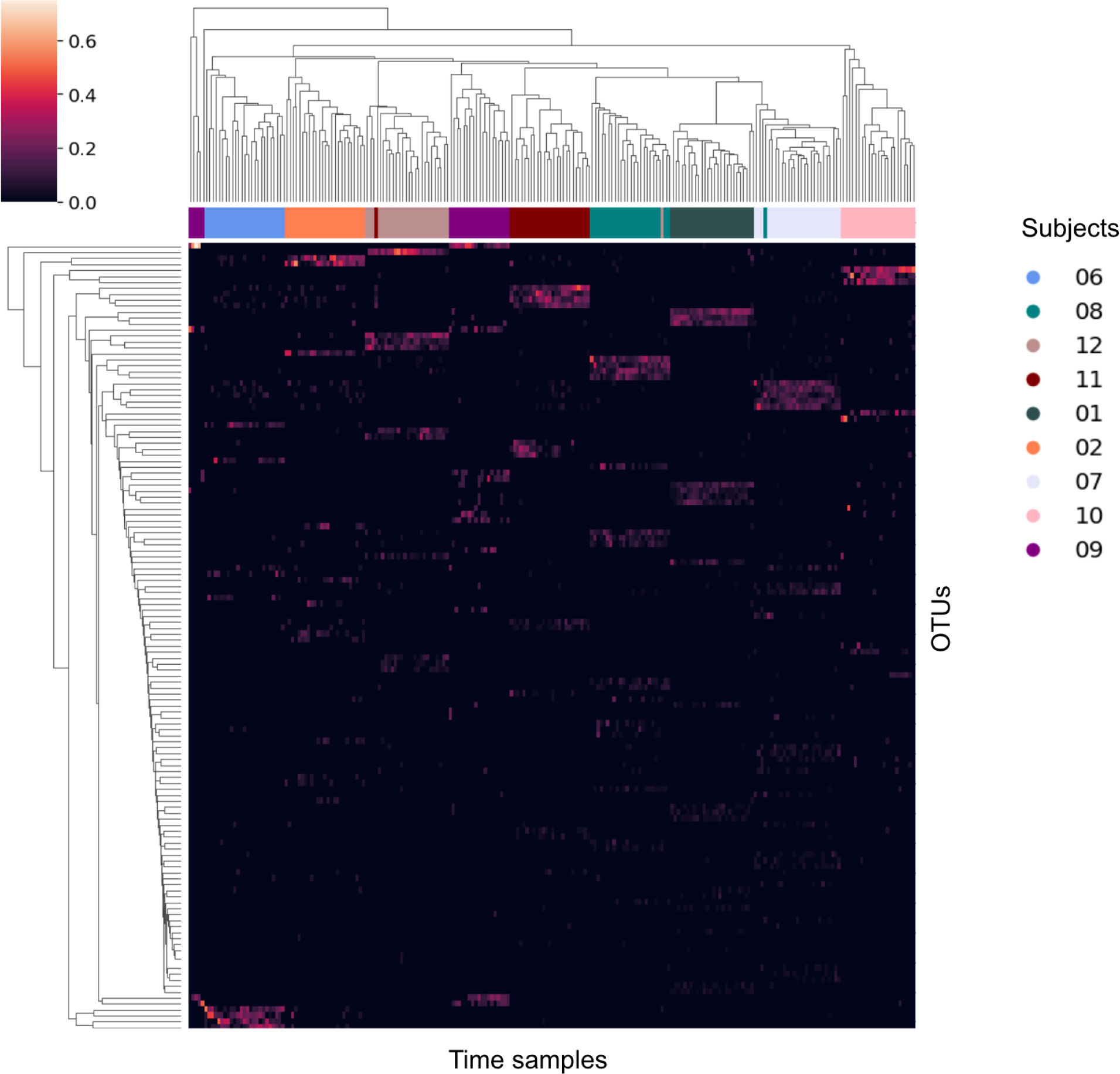
Hierarchical clustering of samples from different subjects based on 226 OTUs which were randomly subsampled from a reduced OTU table. The reduced OTU table did not contain the top 10 most abundant OTUs of each sample, which collectively resulted in the removal of 577 most abundant OTUs. Time samples from each subject are color-coded, and the relative abundance of each OTU is shown in this clustermap.

**SI Figure 13.**
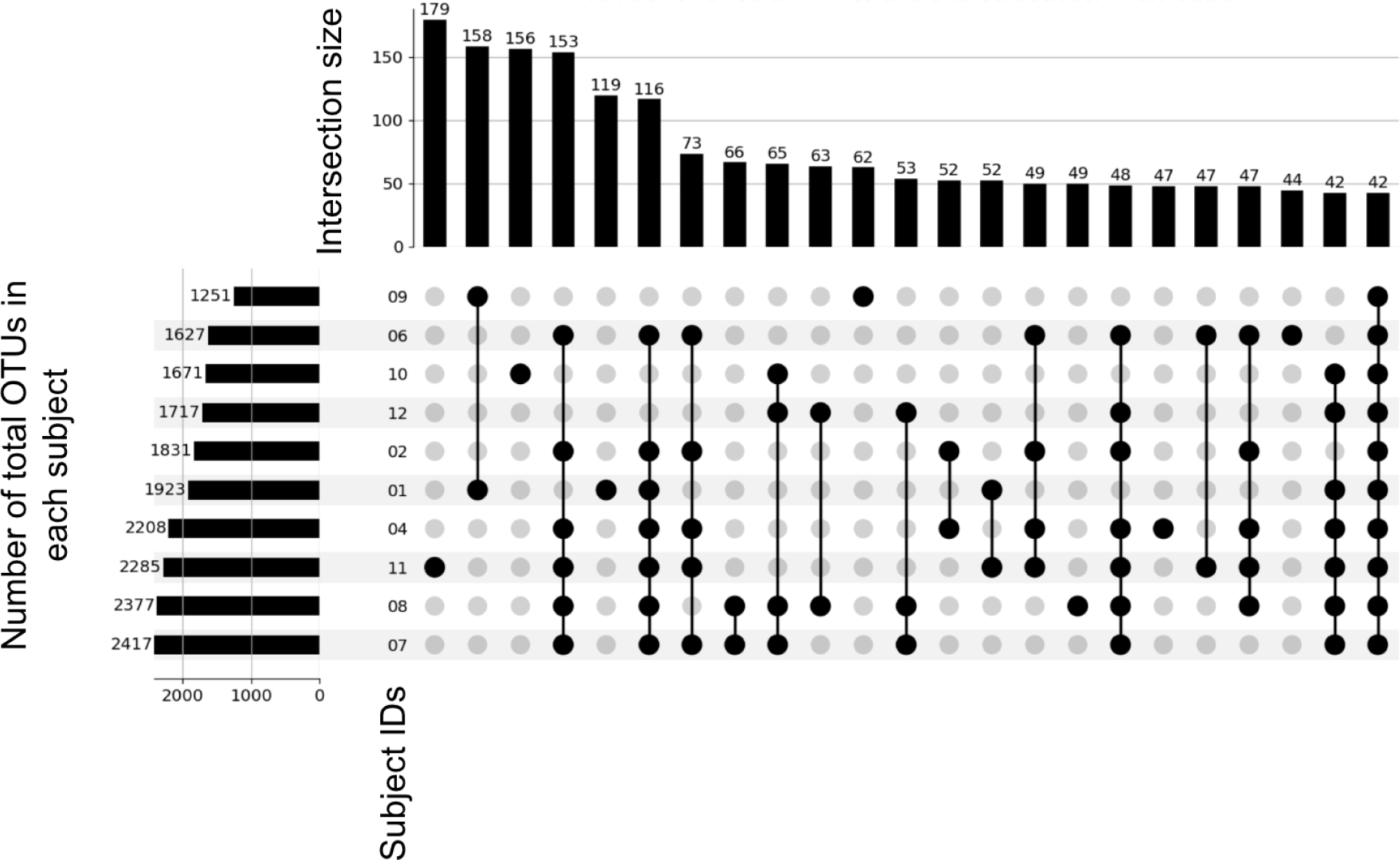
An UpSet plot showing the number of OTUs unique to each subject as well as the number of OTUs shared between different subjects. Connected dots represent sharing of OTUs between one or more subjects, whereas lone dots are sets of OTUs unique to an individual. Sets are ordered based on size (vertical bars). The counts of total OTUs in a subject are shown as horizontal bars for each subject. For visual clarity, sets shown have a minimum size of 40 OTUs.

**SI Figure 14.**
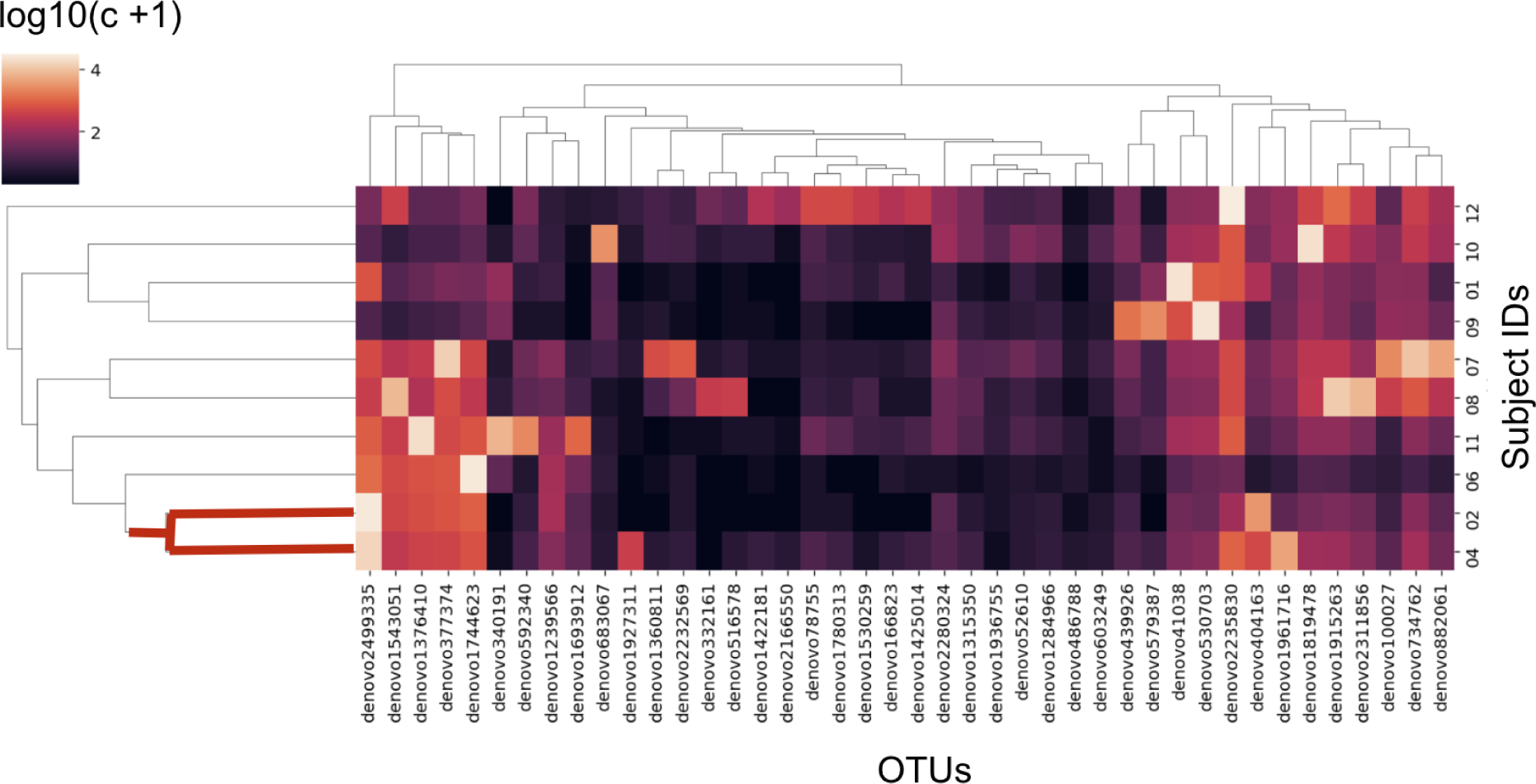
Hierarchical clustering of generalist OTUs-those OTUs found across all subjects within the temporal study-based on their abundance. Counts, c, are based on the abundance of each OTU summed across all time points within a subject. Log10 of counts + 1 is shown in the clustermap. Samples from subjects 2 and 4 who are partners cluster most closely to each other than to other samples.

**SI Figure 15.**
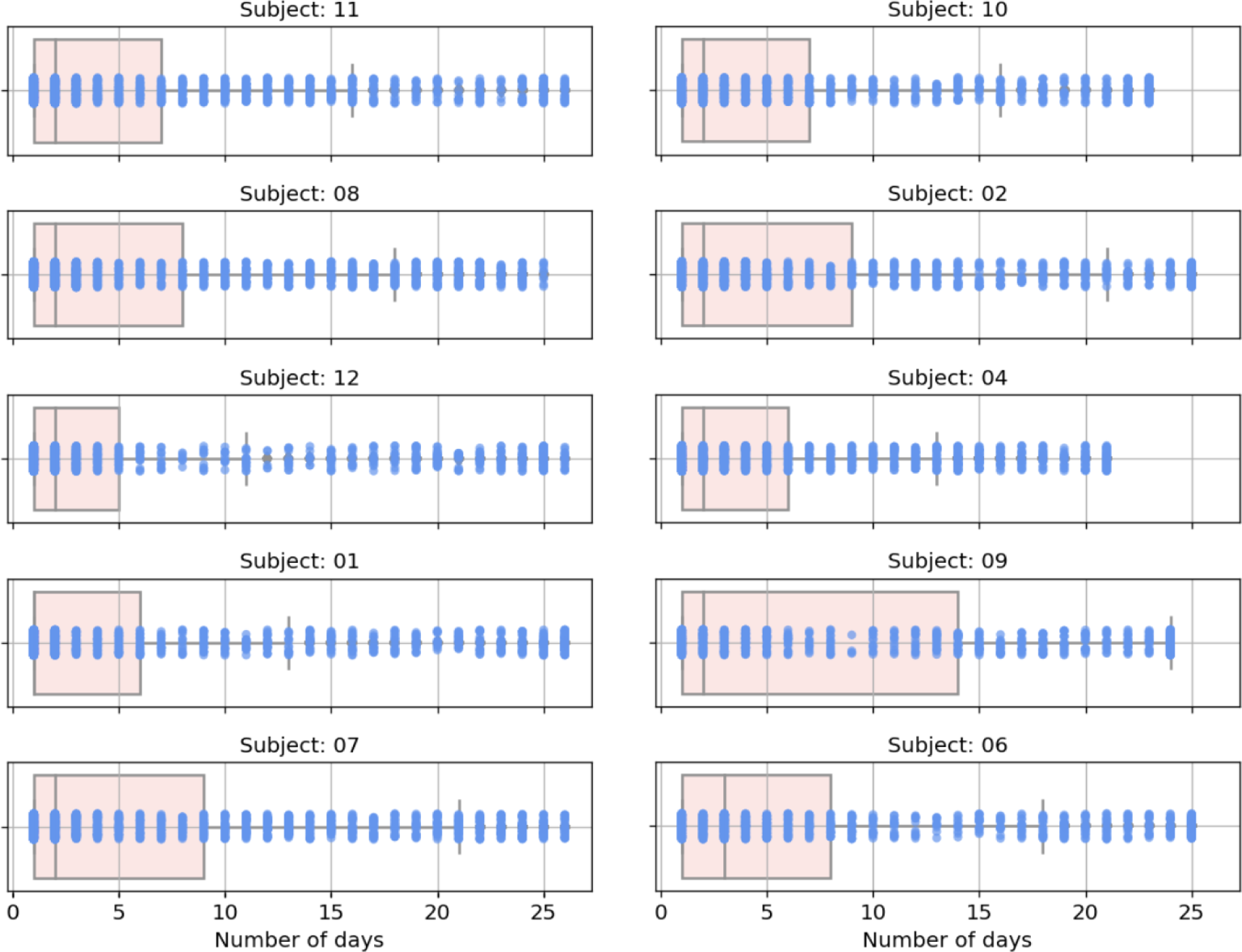
Boxplots and swarmplots of the total number of days that each OTU was observed within an individual. Blue dots correspond to individual OTUs.

**SI Figure 16.**
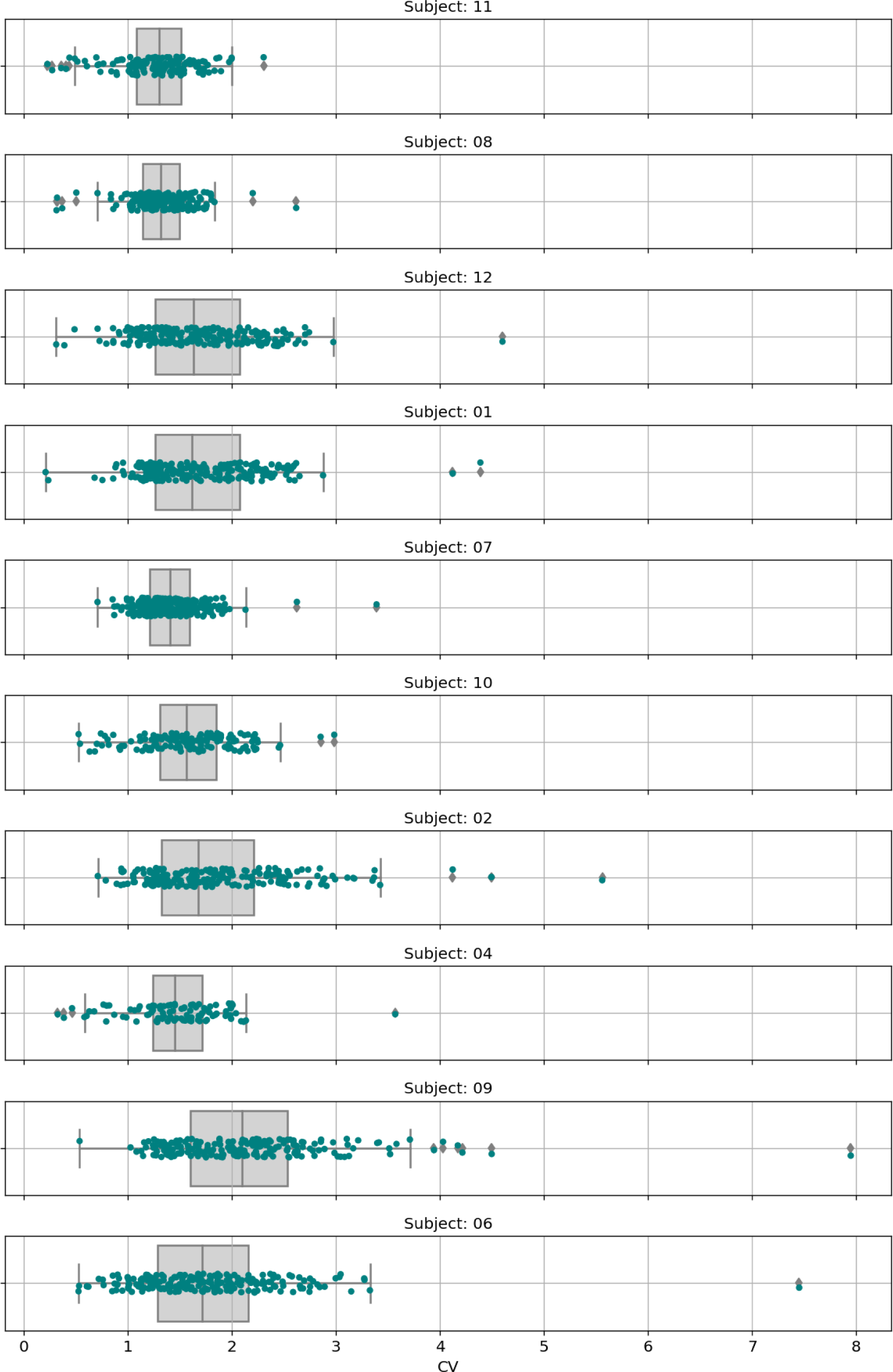
Boxplots and swarmplots showing within each subject the coefficient of variation of persistent OTUs across the sampling period. Persistent OTUs are those that appear in at least 20 days.

**SI Figure 17.**
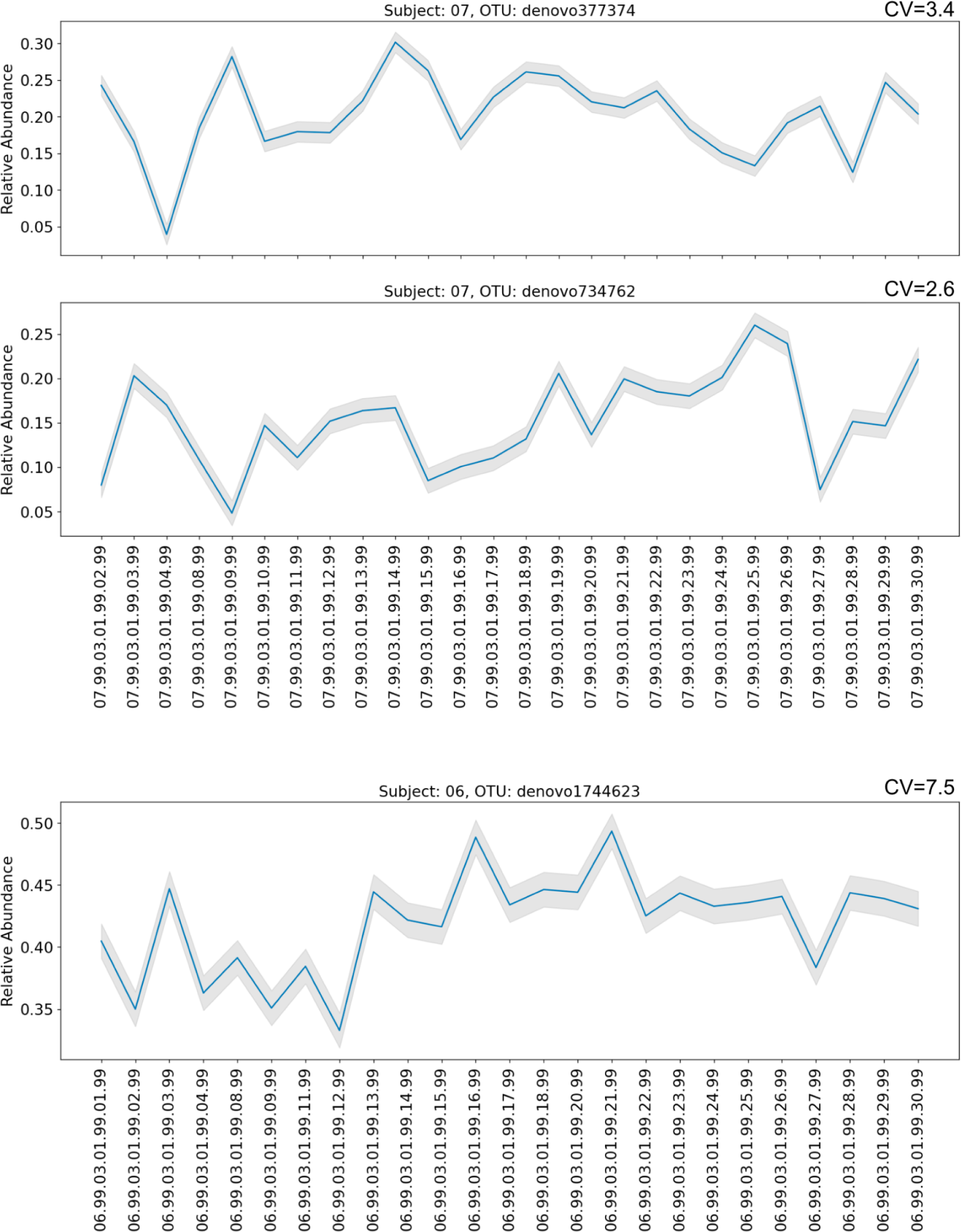
Representative plots of relative abundance as a function of time for outlier OTUs with high coefficient of variation (shown at the top right corners of each plot) in subjects 7 and 6. Gray regions correspond to experimental margin of error. They represent 2 standard deviations above and below the mean. Because only one sample per time point is taken per individual, we conservatively use the maximum standard deviation observed in our noise measurement experiments.

**SI Figure 18.**
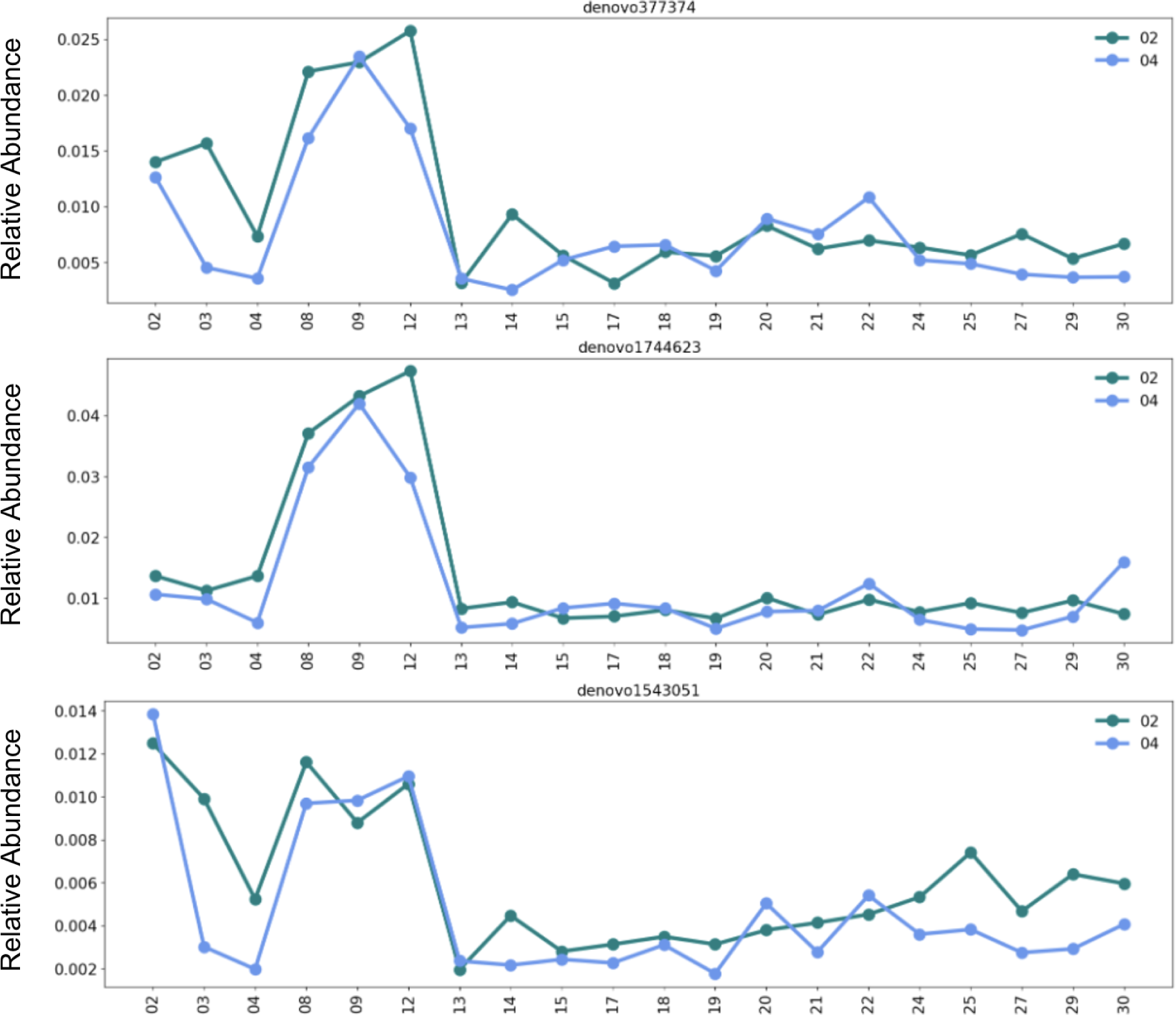
Four OTUs relative abundances shown as a function of time in subjects 2 and 4, who are partners. Only sampling days (x axis) that were made available by both partners are shown for easier comparison.

**SI Figure 19.**
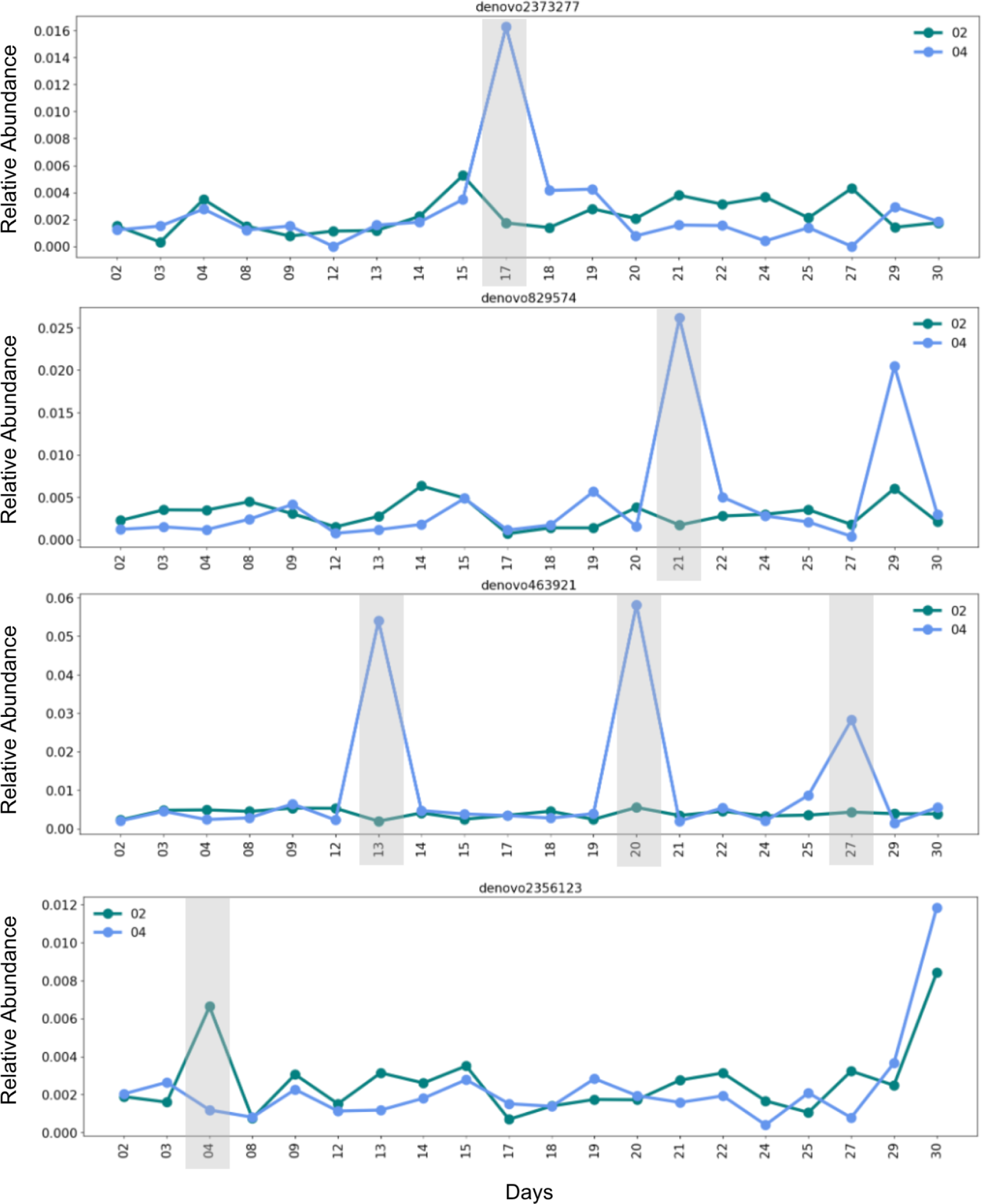
Four OTUs relative abundances shown as a function of time in subjects 2 and 4, who are partners. Grey regions are drawn around time points where there is a several fold change in the relative abundance of a given OTU in one partner but not the other. Only sampling days that were not missed by both partners are shown for easier comparison.

**SI Figure 20.**
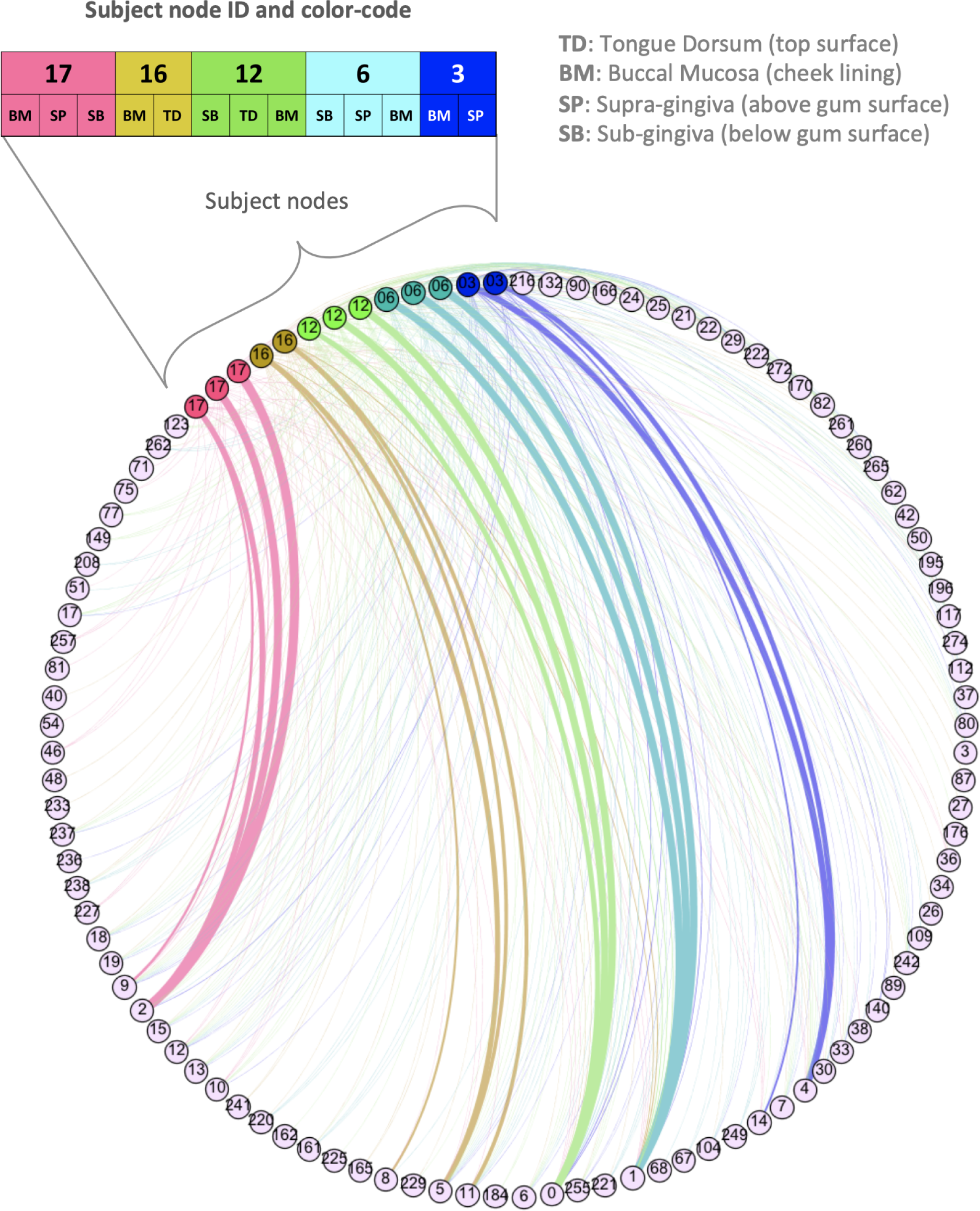
HB1 phage family network. Purple nodes are the OTU nodes and all other nodes represent samples. Sample nodes and edges are color-coded based on the individual they originate from. The oral site associated with each sample is abbreviated next to the sample’s node. Each edge connects an OTU to a sample it exists in, and the edge weight is proportional to the relative abundance of the OTU in that sample. Node IDs are displayed. For OTU nodes, the node ID is the OTU ID. For sample nodes, the nodes IDs are simply the subjects’ IDs.

**SI Figure 21.**
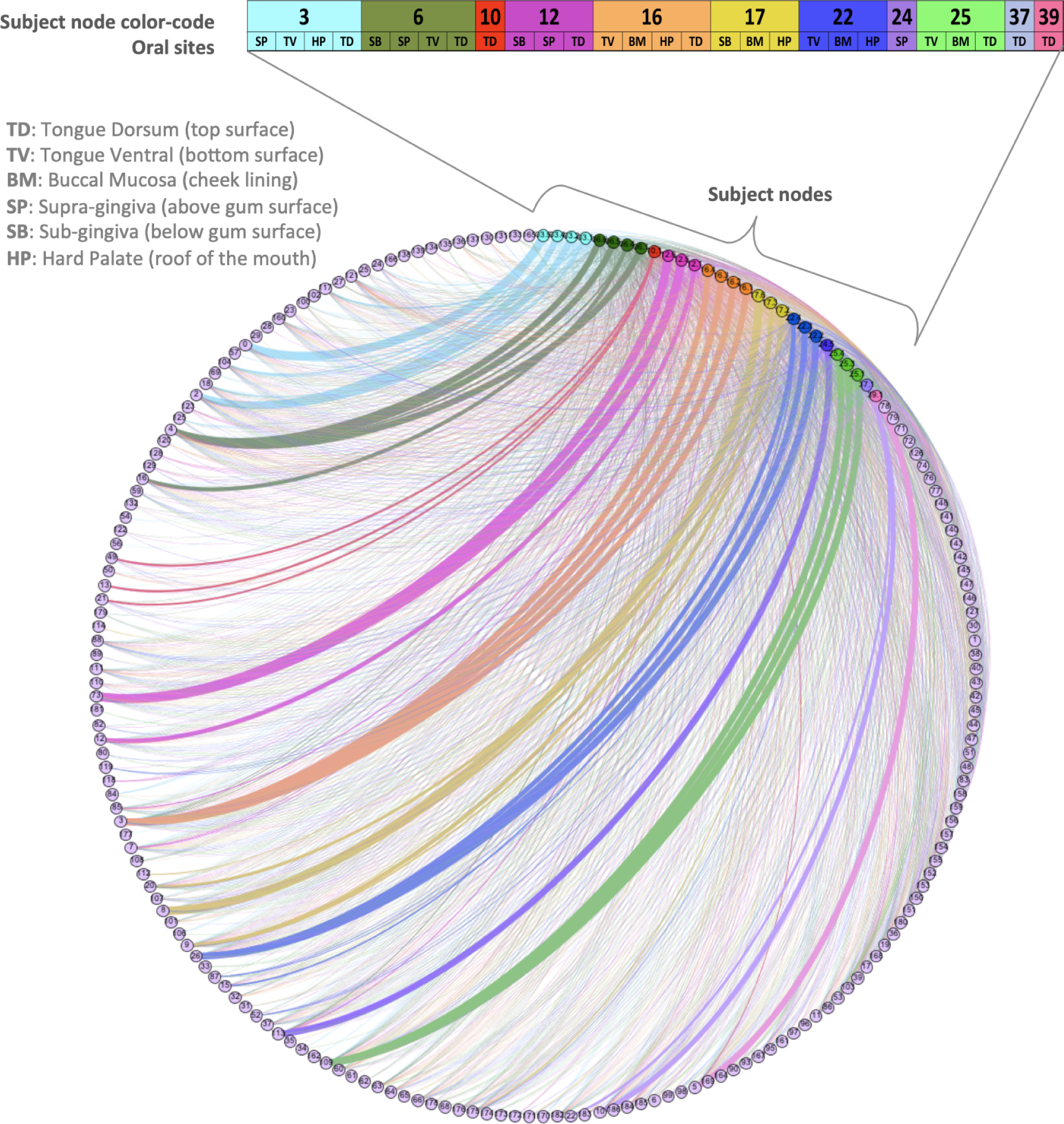
HA phage-host network. Purple nodes are the OTU nodes and all other nodes represent samples. Sample nodes and edges are color-coded based on the individual they originate from. Subject node color code, ID, and the oral sites are displayed above sample nodes. Each edge connects an OTU to a sample it exists in, and the edge weight is proportional to the relative abundance of the OTU in that sample. Node IDs are displayed. For OTU nodes, the node ID is the OTU ID. For sample nodes, the nodes IDs are simply the subjects’ IDs.

**SI Figure 22.**
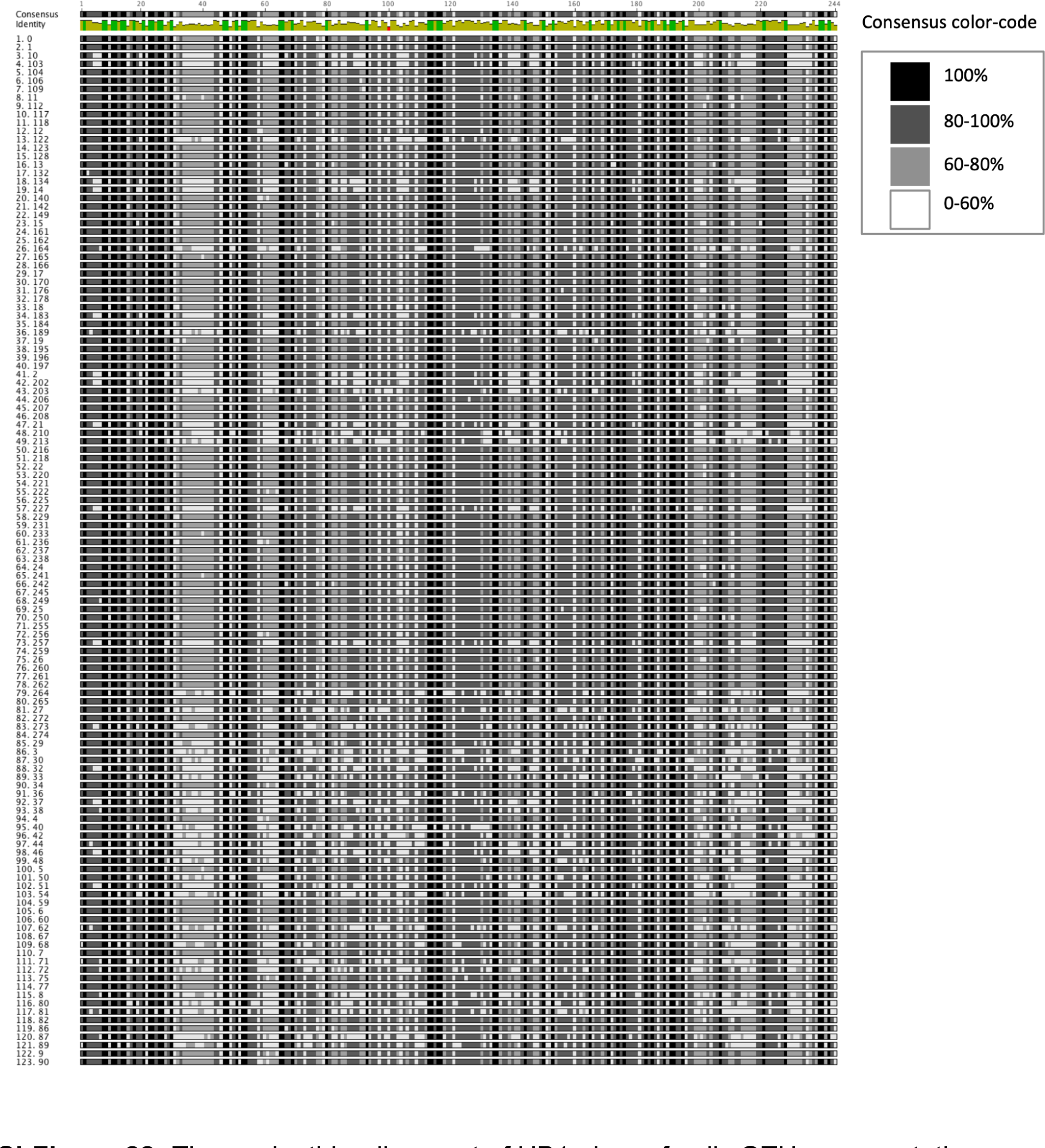
The nucleotide alignment of HB1 phage family OTU representative sequences. Sequences were aligned using Geneious. No gaps were introduced. Each base is color-coded according to its relative abundance within a column in the alignment. Conserved bases are black and highly variable sites are shown in white.

**SI Table 1.**
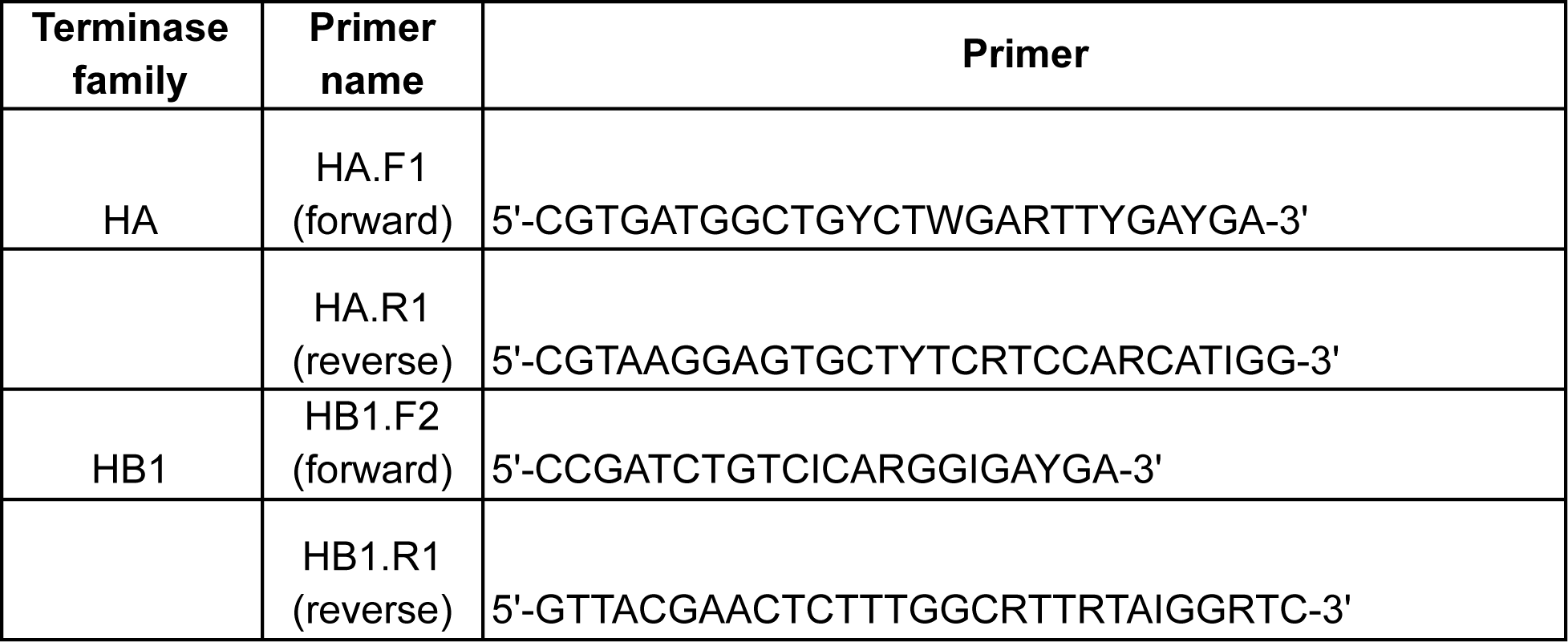
HA and HB1 primer sequences. Please refer to our earlier manuscript for more information about primer design^35^.

**SI Table 2.** Four types of machine learning models are built to classify one individual’s phageprints from the rest (for HB1 terminase family). For each model type, 10 independent models based on 10 different train/test splits are built, and the 95% confidence interval for the Area Under the Precision Recall (AUPR) curve is reported in this table. For instance, column one shows the performance of all model types for detecting subject 6’s phageprint from the rest.

|  | 06 | 08 | 12 | 11 | 01 | 02 | 07 | 10 | 09 |
| --- | --- | --- | --- | --- | --- | --- | --- | --- | --- |
| model |  |  |  |  |  |  |  |  |  |
| **GBC** | (1.0, 1.0) | (0.82, 0.98) | (0.91, 0.99) | (1.0, 1.0) | (1.0, 1.0) | (1.0, 1.0) | (0.92, 1.0) | (1.0, 1.0) | (1.0, 1.0) |
| **KNN** | (1.0, 1.0) | (0.96, 1.0) | (0.96, 1.0) | (0.97, 1.0) | (0.93, 1.0) | (0.96, 1.01) | (0.99, 1.01) | (1.0, 1.0) | (1.0, 1.0) |
| **LR** | (1.0, 1.0) | (0.83, 1.03) | (0.97, 1.01) | (1.0, 1.0) | (1.0, 1.0) | (1.0, 1.0) | (0.99, 1.0) | (1.0, 1.0) | (1.0, 1.0) |
| **MLP** | (1.0, 1.0) | (0.87, 1.02) | (0.96, 1.0) | (1.0, 1.0) | (1.0, 1.0) | (1.0, 1.0) | (0.99, 1.0) | (1.0, 1.0) | (1.0, 1.0) |

**SI Table 3.**
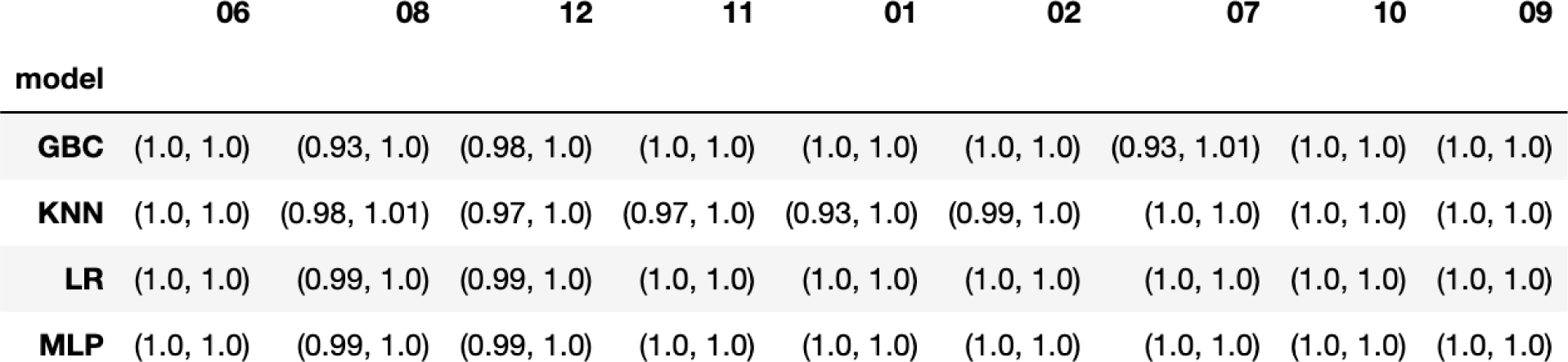
Four types of machine learning models are built to classify one individual’s phageprints from the rest (for HB1 terminase family). For each model type, 10 independent models based on 10 different train/test splits are built, and the 95% confidence interval for the Area Under the Receiver Operator curve (AUROC) is reported in this table. For instance, column one shows the performance of all model types for detecting subject 6’s phageprint from the rest.

**SI Table 4.**
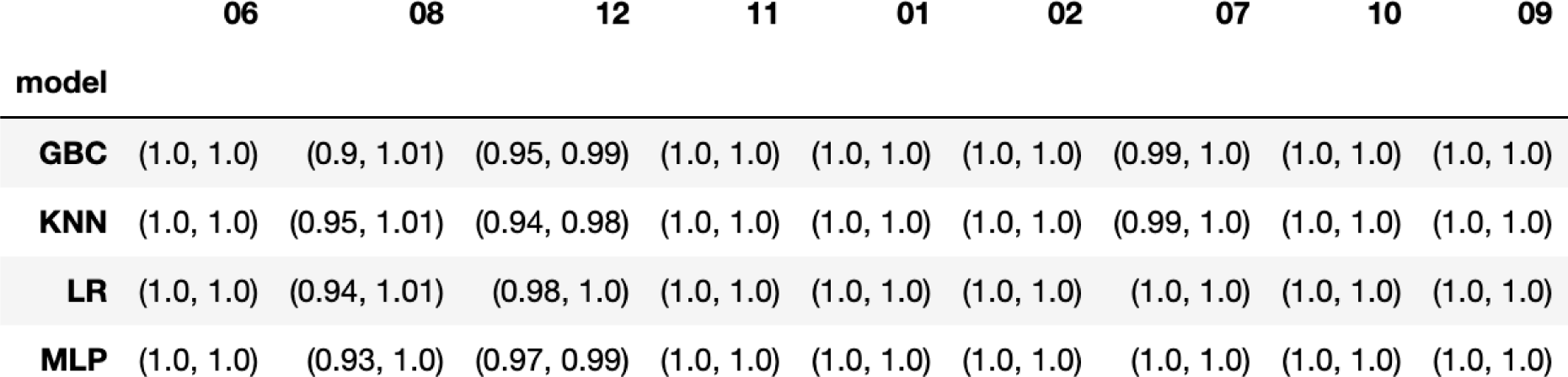
Four types of machine learning models are built to classify one individual’s phageprints from the rest (for HA terminase family). For each model type, 10 independent models based on 10 different train/test splits are built, and the 95% confidence interval for the Area Under the Receiver Operator curve is reported in this table.

**SI Table 5.**
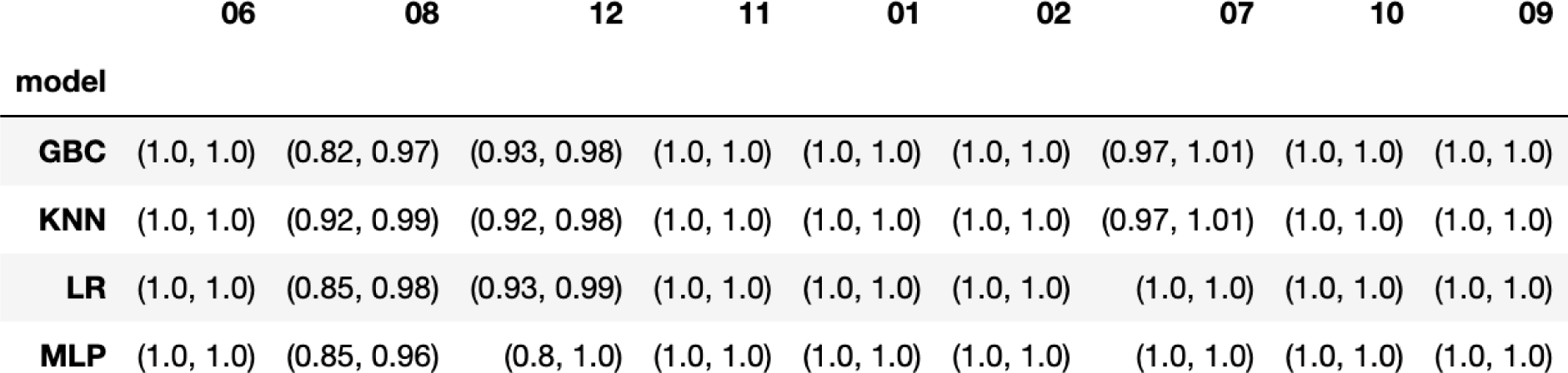
Four types of machine learning models are built to classify one individual’s phageprints from the rest (for HA terminase family). For each model type, 10 independent models based on 10 different train/test splits are built, and the 95% confidence interval for the Area Under the Precision Recall curve is reported in this table.

|  | 06 | 08 | 12 | 11 | 01 | 02 | 07 | 10 | 09 |
| --- | --- | --- | --- | --- | --- | --- | --- | --- | --- |
| model |  |  |  |  |  |  |  |  |  |
| **GBC** | (1.0, 1.0) | (0.82, 0.97) | (0.93, 0.98) | (1.0, 1.0) | (1.0, 1.0) | (1.0, 1.0) | (0.97, 1.01) | (1.0, 1.0) | (1.0, 1.0) |
| **KNN** | (1.0, 1.0) | (0.92, 0.99) | (0.92, 0.98) | (1.0, 1.0) | (1.0, 1.0) | (1.0, 1.0) | (0.97, 1.01) | (1.0, 1.0) | (1.0, 1.0) |
| **LR** | (1.0, 1.0) | (0.85, 0.98) | (0.93, 0.99) | (1.0, 1.0) | (1.0, 1.0) | (1.0, 1.0) | (1.0, 1.0) | (1.0, 1.0) | (1.0, 1.0) |
| **MLP** | (1.0, 1.0) | (0.85, 0.96) | (0.8, 1.0) | (1.0, 1.0) | (1.0, 1.0) | (1.0, 1.0) | (1.0, 1.0) | (1.0, 1.0) | (1.0, 1.0) |

**SI Table 6.** Logistic regression one-versus-rest models built to classify one individual’s HB1 phageprints after ∼600 dominant OTUs have been removed from the dataset. For each model type, 10 independent models based on 10 different train/test splits are built, and the 95% confidence intervals, mean and standard deviation for the Area Under the Receiver Operator Curve (AUROC) is reported in this table.

|  | subject | AUROC_95perc_conf_int | Mean | Std |
| --- | --- | --- | --- | --- |
| **0** | 06 | (1.0, 1.0) | 1.000 | 0.000000 |
| **1** | 08 | (0.99, 1.0) | 0.997 | 0.004830 |
| **2** | 12 | (0.99, 1.0) | 0.998 | 0.004216 |
| **3** | 11 | (1.0, 1.0) | 1.000 | 0.000000 |
| **4** | 01 | (1.0, 1.0) | 1.000 | 0.000000 |
| **5** | 02 | (1.0, 1.0) | 1.000 | 0.000000 |
| **6** | 07 | (1.0, 1.0) | 1.000 | 0.000000 |
| **7** | 10 | (1.0, 1.0) | 1.000 | 0.000000 |
| **8** | 09 | (1.0, 1.0) | 1.000 | 0.000000 |

**SI Table 7.**
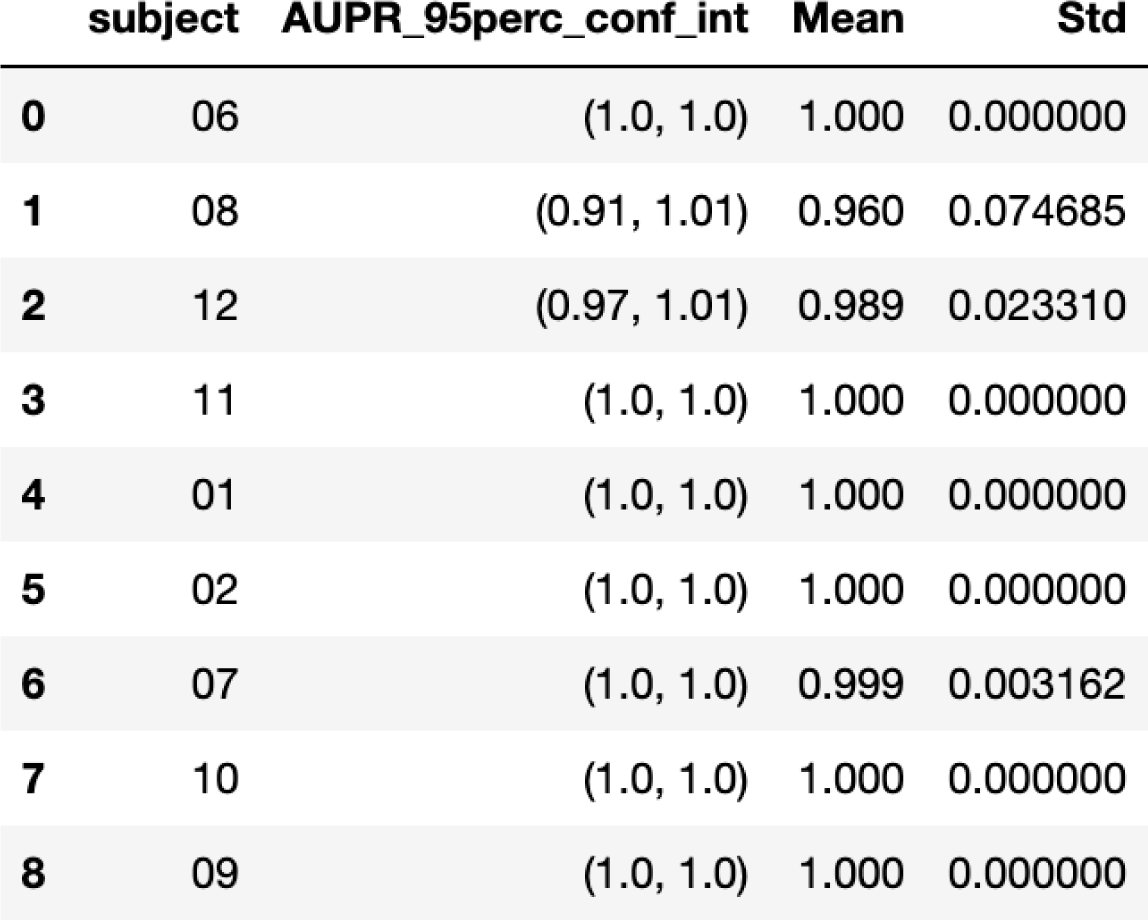
Logistic regression one-versus-rest models built to classify one individual’s HB1 phageprints after ∼600 dominant OTUs have been removed from the dataset. For each model type, 10 independent models based on 10 different train/test splits are built, and the 95% confidence intervals, mean and standard deviation for the Area Under the Precision Recall curve (AUPR) is reported in this table.

**SI Table 8.**
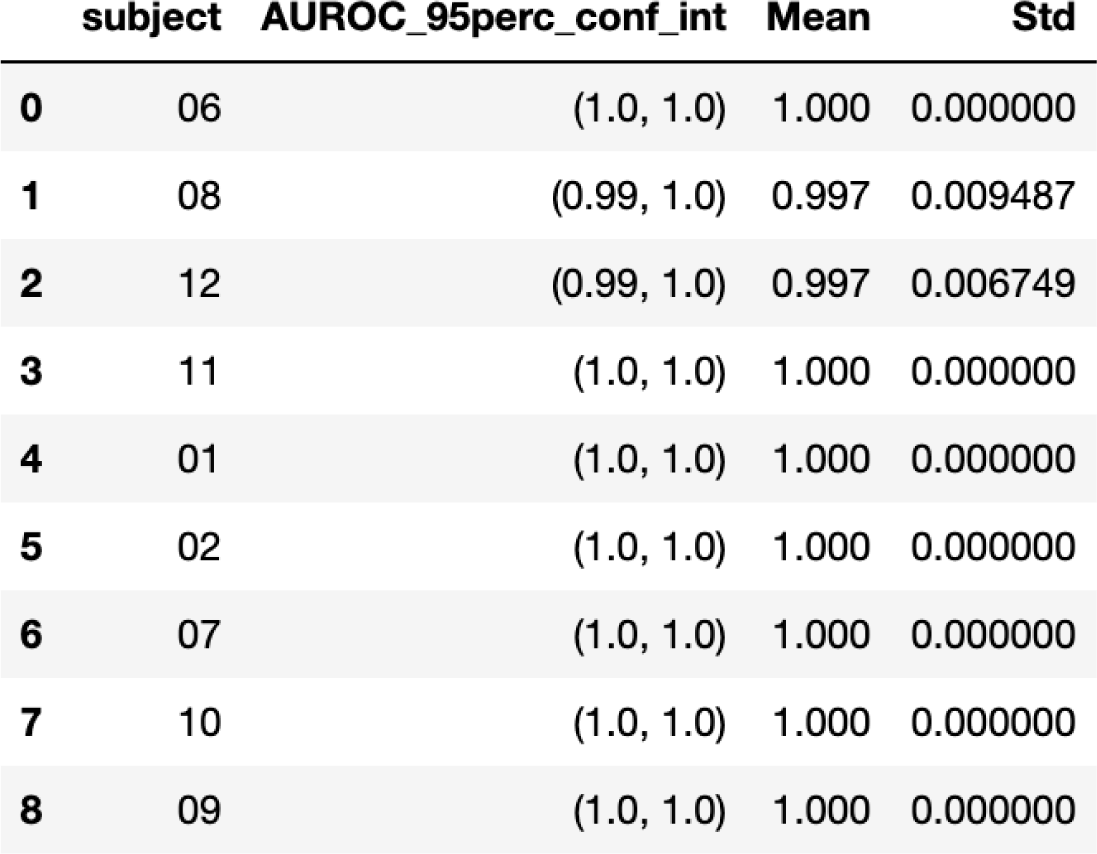
Logistic regression one-versus-rest models built to classify one individual’s HB1 phageprints after ∼600 dominant OTUs were removed and the resulting OTU table was subsampled to contain only 2% of the total OTUs (∼200). For each model type, 10 independent models based on 10 different train/test splits are built, and the 95% confidence intervals, mean and standard deviation for Area Under the Receiver Operator Curve (AUROC) is reported in this table.

|  | subject | AUROC_95perc_conf_int | Mean | Std |
| --- | --- | --- | --- | --- |
| **0** | 06 | (1.0, 1.0) | 1.000 | 0.000000 |
| **1** | 08 | (0.99, 1.0) | 0.997 | 0.009487 |
| **2** | 12 | (0.99, 1.0) | 0.997 | 0.006749 |
| **3** | 11 | (1.0, 1.0) | 1.000 | 0.000000 |
| **4** | 01 | (1.0, 1.0) | 1.000 | 0.000000 |
| **5** | 02 | (1.0, 1.0) | 1.000 | 0.000000 |
| **6** | 07 | (1.0, 1.0) | 1.000 | 0.000000 |
| **7** | 10 | (1.0, 1.0) | 1.000 | 0.000000 |
| **8** | 09 | (1.0, 1.0) | 1.000 | 0.000000 |

**SI Table 9.** Logistic regression one-versus-rest models built to classify one individual’s HB1 phageprints after ∼600 dominant OTUs were removed and the resulting OTU table was subsampled to contain only 2% of the total OTUs (∼200). For each model type, 10 independent models based on 10 different train/test splits are built, and the 95% confidence intervals, mean and standard deviation for the Area Under the Precision Recall curve (AUPR) is reported in this table.

|  | subject | AUPR_95perc_conf_int | Mean | Std |
| --- | --- | --- | --- | --- |
| 0 | 06 | (1.0, 1.0) | 1.000 | 0.000000 |
| 1 | 08 | (0.96, 1.01) | 0.984 | 0.032042 |
| 2 | 12 | (0.97, 1.01) | 0.987 | 0.027508 |
| 3 | 11 | (1.0, 1.0) | 1.000 | 0.000000 |
| 4 | 01 | (1.0, 1.0) | 1.000 | 0.000000 |
| 5 | 02 | (1.0, 1.0) | 0.999 | 0.003162 |
| 6 | 07 | (1.0, 1.0) | 0.999 | 0.003162 |
| 7 | 10 | (1.0, 1.0) | 1.000 | 0.000000 |
| 8 | 09 | (1.0, 1.0) | 1.000 | 0.000000 |

**SI Table 10.**
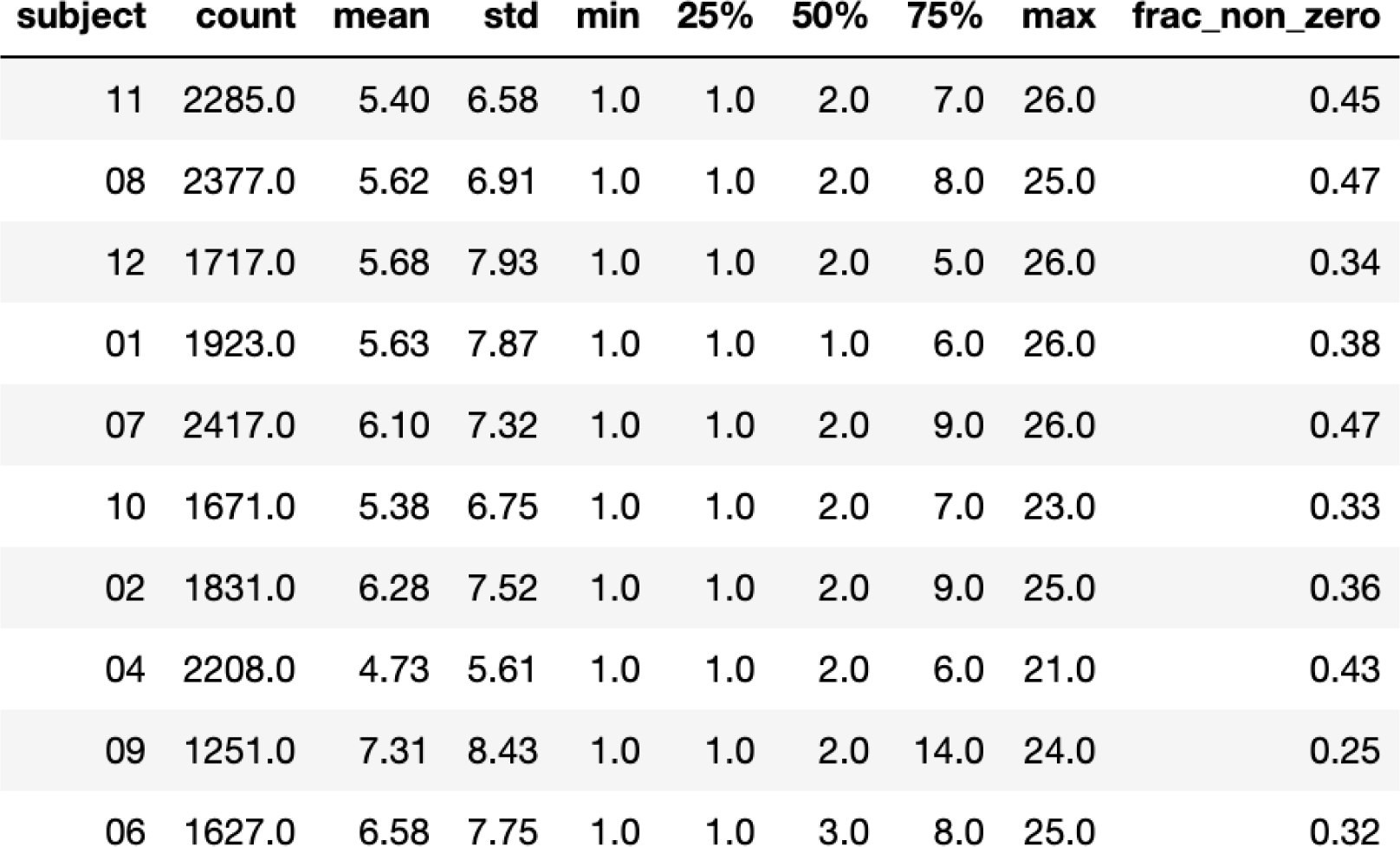
Summary statistics for SI Figure 15. The absolute count and fraction of OTUs detected at least once in an individual are reported under the “count” and “frac_non_zero” columns, respectively. The mean, standard deviation, and median number of days OTUs are detected among other statistics are reported.

## Notes

### Competing Interest Statement

The authors have declared no competing interest.

